# Traction force reconstruction assessment on real three-dimensional matrices and cellular morphologies

**DOI:** 10.1101/2022.11.16.516745

**Authors:** Alejandro Apolinar-Fernández, Jorge Barrasa-Fano, Mar Cóndor, Hans Van Oosterwyck, José A. Sanz-Herrera

**Affiliations:** Escuela Técnica Superior de Ingeniería, Universidad de Sevilla; Biomechanics section, Department of Mechanical Engineering, KU Leuven; Prometheus division of Skeletal Tissue Engineering, KU Leuven

**Keywords:** Traction Force Microscopy, Mechanobiology, Nonlinear mechanics, Computational mechanics, Finite element method

## Abstract

Traction force microscopy (TFM) allows to estimate tractions on the surface of cells when they mechanically interact with hydrogel substrates that mimic the extracellular matrix (ECM). The field of mechanobiology has a strong interest in using TFM in 3D in vitro models. However, there are a number of challenges that hamper the accuracy of 3D TFM and that are often bypassed. In this study, the computational efficiency and accuracy of TFM, referred to traction reconstruction from synthetically generated (control) ground truth solutions, are assessed from four different perspectives: magnitude of cellular pulling force (and hence strain level achieved in the hydrogel), effect of the complexity of the cellular morphology, accuracy and computational efficiency of forward vs inverse traction recovery methods, and the effect of incorrectly selecting a constitutive model that describes the behavior of the ECM (i.e. linear/nonlinear). The main results showed: (i) traction reconstruction is more challenging for complex cell morphologies, (ii) there is no significant impact of the magnitude of cellular pulling force on the overall reconstruction accuracy, and (iii) modeling a nonlinear hydrogel with a linear constitutive model leads to non-negligible errors (up to 80% and 30% for forward and inverse methodologies, respectively) in traction reconstruction. This study expands the characterization of the accuracy and efficiency of 3D TFM, highlighting important factors to be considered in future 3D TFM in vitro applications.

## 1. Introduction

The mechanical interactions of cells with the extracellular matrix (ECM) are crucial for multiple physiological and pathological processes (Ingber, 2003; Mammoto et al., 2013; Vogel and Sheetz, 2006). Over the last decades, the field of mechanobiology has discovered that cell-ECM mechanical interactions can drive processes such as cancer invasion (Jiang et al., 2019; Kopanska et al., 2016; Kumar and Weaver, 2009; Medina et al., 2019; Nguyen-Ngoc et al., 2012; Peng et al., 2019; Shields et al., 2007), angiogenesis (Elliott et al., 2015; Ingber, 2002; Vaeyens et al., 2020) or stem cell differentiation (Discher et al., 2005; Engler et al., 2006; Guilak et al., 2009; Huebsch et al., 2010). In vitro models are typically used to obtain fundamental understanding of cellular mechanotransduction pathways that can become pivotal in developing tissue engineering and therapeutic techniques (Janmey et al., 2020; Jansen et al., 2015; Kim et al., 2021; Kutys and Chen, 2016; Vining and Mooney, 2017).

In order to extract information from these systems, researchers make use of optical microscopy combined with computational methods that include microscopy image analysis techniques and mechanical models. One such methodology is Traction Force Microscopy (TFM), which has been the main choice for calculating forces exerted by cells on the ECM for the last decades (Butler et al., 2002; Dembo and Wang, 1999; Harris et al., 1980; Sabass et al., 2008). In TFM, cells are seeded on top of or embedded in a substrate or hydrogel that contains fiducial markers (often fluorescent beads) and that mimics the ECM. Microscopy images are acquired before (with mechanically active cells; stressed state) and after (relaxed state) cell removal or cell mechanical inhibition. The matrix deformations caused by cellular forces lead to changes in the position of the markers, which can be measured by means of image analysis algorithms that compare the stressed state to the relaxed state images. Finally, a constitutive model that describes the mechanical behavior of the substrate/hydrogel is used to calculate strain, stress and traction forces from the measured displacement field. Several ways of calculating traction forces from the measured displacements have been proposed in the literature. On the one hand, *forward* (also found in the literature as *direct*) methods calculate the strain field by differentiating the measured displacements, and the stress/traction field is computed by means of the constitutive relation for the ECM (Franck et al., 2011; Gjorevski et al., 2015; Toyjanova et al., 2014). On the other hand, *inverse* methods search for a traction solution that is consistent with the measured displacements while being subject to a certain constraint that limits the effect of measurement noise in the resultant traction field (Feng and Hui, 2016; Legant et al., 2010).

TFM is very well established for 2D in vitro cultures where cells are seeded on top of a substrate. Polyacrylamide substrates are typically chosen due to their linear elastic properties. This allows for using simple analytical formulations such as the Boussinesq solution, which heavily eases traction recovery (Huang et al., 2020; Izquierdo-Álvarez et al., 2019). However, the dimensionality of the surrounding microenvironment is crucial for cell behavior and the field considers 3D in vitro cultures as more physiologically relevant (Duval et al., 2017; Vogel and Sheetz, 2006). As a result, researchers have started using other bio-mimetic materials that allow for embedding cells in a 3D environment. Synthetic materials of highly tunable properties such as Polyethylene Glycol (PEG) as well as natural materials that more closely resemble physiological conditions such as collagen or fibrin hydrogels have been primarily used (Caliari and Burdick, 2016). This has lead to the development of more complex 3D TFM methodologies in the last twelve years.

The approach chosen to recover cellular tractions will vary depending on two factors. First, the stress-strain relationship of the hydrogel can be linear or nonlinear. While PEG hydrogels can be tuned to display a linear constitutive response, fibrillar hydrogels are characterized by nonlinear responses and strain stiffening. Second, researchers often use the small strain theory to describe the behavior of the hydrogel. However, when deformations are large, this theory is no longer valid and the finite-strain theory should be adopted.

The literature offers a vast number of approaches combining the previous two factors differently. 3D traction forces have been calculated in PEG hydrogels using linear elastic models and under a small strain elasticity theory for single cells (Legant et al., 2010) and for angiogenic sprouts (Barrasa-Fano et al., 2021a,b). Collagen hydrogels have been modeled as a linear elastic material (Du et al., 2018; Gjorevski et al., 2015) as well as a hyperelastic material such as Neo-Hookean (Hervas-Raluy et al., 2021; Sanz-Herrera et al., 2021; Song et al., 2020b). A particular case of application in this study is the work of Steinwachs et al. (2016), who developed a specific nonlinear constitutive model that best fit the experimentally acquired (by means of e.g. shear rheology mechanical testing) strain-stress curves of a collagen hydrogel.

However, the choice of a constitutive model or an elasticity theory are often taken arbitrarily either to simplify the study or to highlight its increased complexity with respect to the state of the art. There is a lack of systematic analysis on the effect of incorrectly selecting a model. Recently, Song et al. (2020a) developed a traction recovery algorithm that accounts for both material and geometric nonlinearities. Interestingly, they quantified the effect of neglecting nonlinearity by resolving cellular forces synthetically generated with a nonlinear model (ground truth) with a linear model. However, they only explored this effect for an arbitrary traction field of a given magnitude. Existing studies have not analyzed the effect of the magnitude of the cell tractions on the accuracy of TFM.

Moreover, we previously reported on the severe impact of the size and distribution of focal adhesions on 3D TFM accuracy (Barrasa-Fano et al., 2021a). Indeed, it is important to analyze factors that lead to traction fields that are difficult to recover. Another such factor is the complexity of the geometry of a cell. While initial studies validated their traction recovery algorithm in simplified geometries such as cylinders (Du et al., 2018), spheres (Holenstein et al., 2019) or ellipsoids (Vitale et al., 2012), later studies have used real cell geometries obtained from the segmentation of a microscopy image of a cell to make their results more representative of a real scenario (Barrasa-Fano et al., 2021a; Hervas-Raluy et al., 2021; Sanz-Herrera et al., 2021; Song et al., 2020b). However, these studies always perform simulations on a sole cell geometry. Cell geometries can be highly heterogeneous going from spherical to spread and protrusive. This depends on multitude of factors such as cell type, hydrogel properties (adhesiveness, degradability, stiffness, porosity, etc), or incubation time. There are no studies that systematically report on how traction calculation accuracy is affected by the geometry of a cell.

The aim of this study is to assess the accuracy and efficiency of TFM, with regard to traction reconstruction versus a (controlled) synthetic ground truth solution, from different perspectives: cellular pulling force magnitude and hence strain level achieved in the hydrogel, effect of cellular morphology complexity, accuracy and computational efficiency of forward vs inverse traction reconstruction approaches, and the effect of incorrectly selecting a constitutive model that describes the behavior of the ECM (i.e. linear/nonlinear). The paper is organized as follows: first, the theoretical bases of the methods used in this study (forward and inverse) are briefly introduced. Then, the different cases of study and in silico TFM methodology and simulations are explained in section 3. The corresponding results are detailed in section 4, and discussed in section 5. Finally, the most important conclusions are drawn at the end of the paper.

## 2. Theoretical background

Traction Force Microscopy can be performed by means of two fundamentally distinct methodologies commonly referred to as the *forward* and the *inverse* methods (Schwarz and Soiné, 2015). The forward method computes the cellular tractions and hydrogel stresses directly from the measured displacements, and the assumed constitutive law of the hydrogel. Alternatively, the inverse method uses this displacement field as input data in order to obtain a traction field that meets certain criteria within a minimization problem framework, based on a regularization strategy. The forward method is less computationally demanding than the inverse method, but the latter provides more precise traction field reconstructions (Barrasa-Fano et al., 2021a). The methods that we use in this study are a conventional forward (direct traction computation from measured displacements via the material’s constitutive law), and an inverse formulation which searches for a new displacement field that approximates the measured one, ensuring that force equilibrium within the hydrogel is satisfied (Sanz-Herrera et al., 2021). Both the forward and inverse methods were developed and implemented in the context of both linear elasticity and finite strain hyperelasticity, under a finite element (FE) approach.

### 2.1. Forward method

The forward method is formulated in this section within the framework of finite strain hyperelasticity. Once given the reconstructed displacements field **u**^*m*^ referring to the spatial (Eulerian) description of the medium **x**, the position of a point in the material description (Lagrangian) **X** is given by,

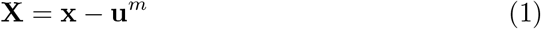

Then, the deformation gradient is defined as,

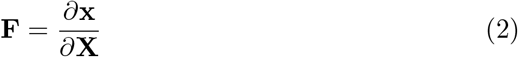

which, by using (1), yields the expression,

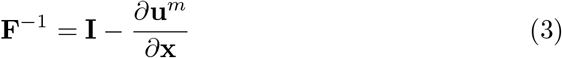

being **I** the identity matrix. It is important to recall that the reference configuration in TFM corresponds to the deformed hydrogel and stressed cell state after which a relaxed (assumed stress free) hydrogel configuration is achieved. After a FE mesh discretization, the displacement field used in this forward method (**u**^*fwd*^) is approximated as follows,

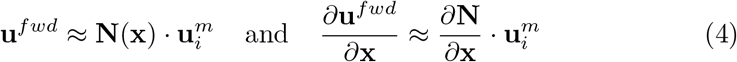

where **N**(**x**) denotes the matrix which contains standard shape functions associated to element interpolation, and the reconstructed displacement field **u**^*m*^ is defined at nodal positions *i*. Then, using Eq. (4) into Eq. (3), the expression for the computation of the discrete inverse of the deformation gradient is given as,

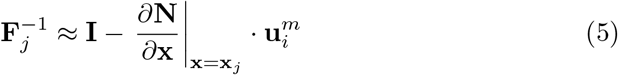

where derivatives are computed at the Gauss points **x**_*j*_ of the finite element mesh.

Once obtained the deformation gradient, stresses are computed from this quantity by use of a certain constitutive relation. The reader is addressed to Appendix A for a detailed description of the nonlinear constitutive law employed in this study and the associated mechanical variables. Finally, tractions are computed following Cauchy’s formula as,

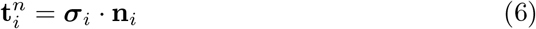

in which 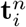 is the traction vector associated to the spatial configuration at nodal points *i* of the finite element mesh, ***σ***_*i*_ is the Cauchy stress tensor at nodal point *i*, once an averaging procedure of *σ*_*j*_ from Gauss points is conducted. Finally, **n**_*i*_ denotes the outward normal vector to node *i*, which is numerically computed at nodal points from the surfaces of elements of the finite element mesh in the deformed (reference) configuration.

### 2.2. Inverse method

The inverse method used in this work, PBNIM (physics-based nonlinear inverse method) (Sanz-Herrera et al., 2021), focuses on finding a new displacement field **u**^*inv*^ that approximates as closely as possible the measured one **u**^*m*^ and that fulfills the equilibrium condition within the medium of interest, i.e., the hydrogel in which the cell is embedded. This can be modeled as a minimization problem that includes the mechanical equilibrium constraint equation, and is mathematically described as follows,

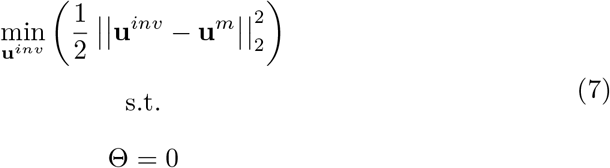

where Θ generically represents the equilibrium constraint manifold where the candidate solution must lie. For the sake of compatibility with its finite element formulation, the functional Θ is developed in terms of the Principle of Virtual Work (PVW) equation. The reader is referred to Appendix A where this elaboration is explained in more detail in the context of its integration with ABAQUS software (used in the numerical implementation) and the chosen constitutive law.

The equilibrium constraint can be integrated into the cost function (7) by means of a continuum and scalar Lagrange multiplier *η* as follows,

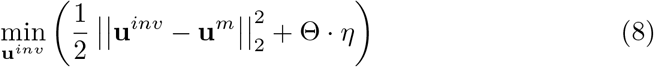

Using the Gateaux derivative, (8) has its analytical minimal stationary solution at,

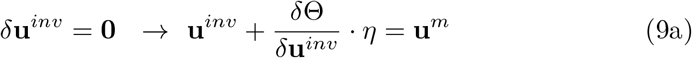

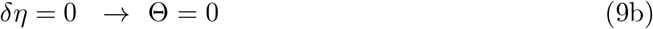

Eqs. (9a) and (9b) are then elaborated, linearized and discretized in a FE framework. The reader is referred to Sanz-Herrera et al. (2021) for further information and details about PBNIM numerical implementation.

## 3. In silico TFM simulations

The study presented in this article follows a holistic approach, which includes a compilation of cases up to a total of 744 in silico simulations. They are summarized in Fig. 1 and are explained next.

**Figure 1:**
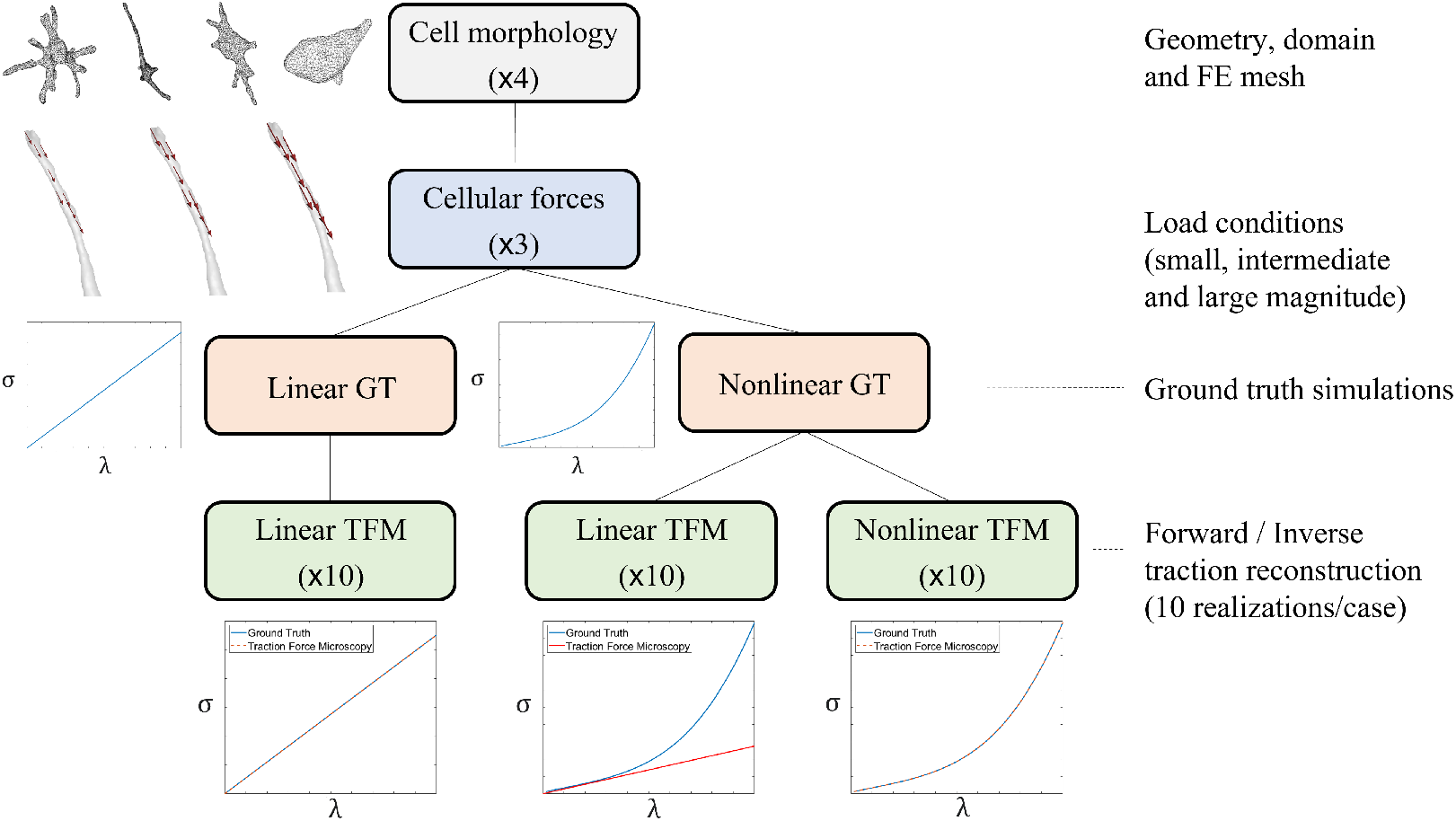
Tree diagram showing the different series of cases considered in the study.

### 3.1. Ground truth cases

First, a number of displacement fields were synthetically reproduced for different cell morphologies, levels of cellular traction magnitudes and different assumed mechanical behaviors of the hydrogels. These displacement fields were directly computed from FE simulations using the software ABAQUS.

#### Cell morphologies

We selected four different single cell morphologies obtained from real confocal microscopy images (voxel size: 0.57×0.57×1 *µm*^3^). Each cell displayed a different degree of morphological complexity. They were classified into 4 categories: (i) Spherical cells, with similar longitudinal and transversal lengths and few protrusions; (ii) Spread cells, whose longitudinal dimension is notably longer than its transversal one; (iii) Protrusive cells, which presents an elongated morphology with protrusions; and (iv) Star-shaped cells, with no apparent dominant dimension in length, but with a large number of protrusions.

Cell morphologies, as well as in silico models, domains of analysis, and loads are shown in Fig. 2. Finite element meshes were created with the Matlab toolbox TFMLAB (Barrasa-Fano et al., 2021b). Moreover, the solidity index, defined as the ratio between the cell volume and the volume of its convex hull, is used as an indicator of cell morphology complexity. Higher solidity indicates less complex (closer to a spherical shape) cell geometries. The solidity value of each cell is plotted in Fig. 3.

**Figure 2:**
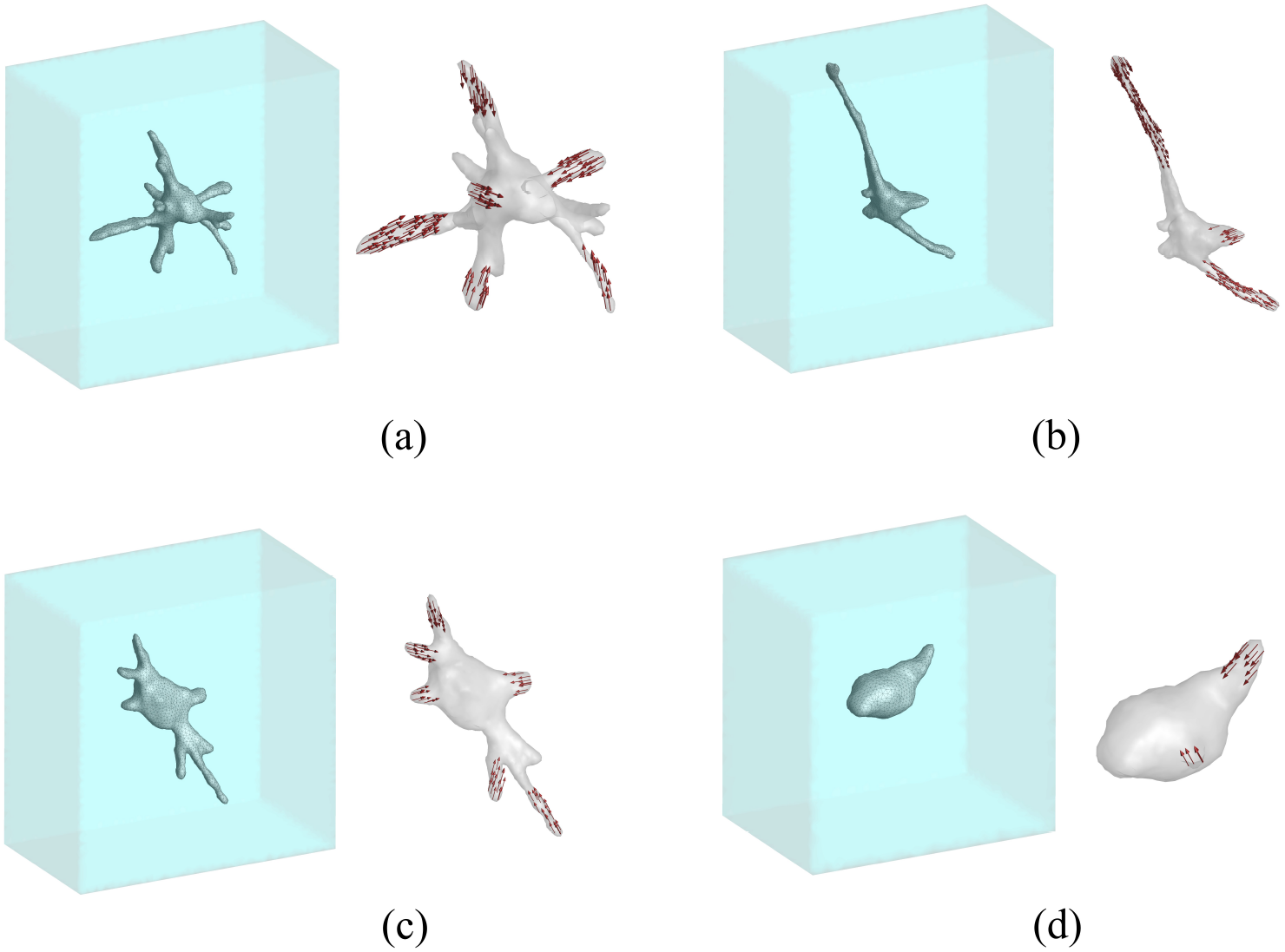
In silico models with selected cell morphologies: (a) star-shaped cell, (b) spread cell, (c) protrusive cell, and (d) spherical cell. The models illustrate the 3D cell embedded in the hydrogel (left part of each subfigure) and the body of the cells including location and direction of prescribed pulling forces (right part of each subfigure).

**Figure 3:**
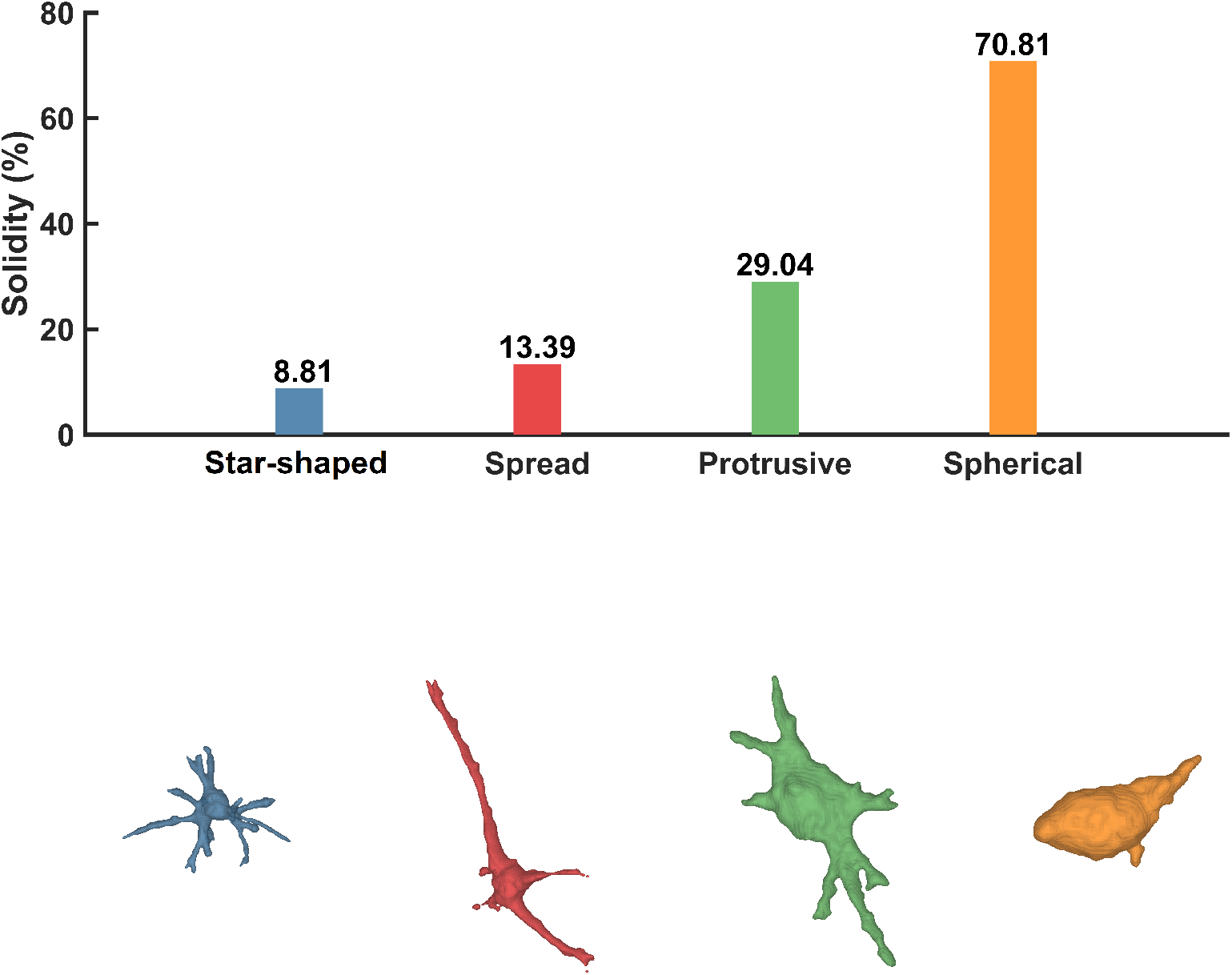
Solidity index for selected cell morphologies.

#### Prescribed cellular tractions

Cellular tractions are prescribed as distributed and uniform nodal forces along the tips of selected protrusions of the different analyzed cells. The direction of the forces is defined along the main axis of the protrusion, and inwardly towards the center of the cell. Fig. 2 shows the regions of application and direction of nodal forces for the different cells.

Three different cellular pulling force magnitudes (small, intermediate and large) were selected in order to investigate their impact on traction reconstruction accuracy. Large cellular pulling force magnitudes were defined such that their associated maximum strains (Frobenius norm of the logarithmic strain tensor, i.e., the Euclidean norm of the logarithmic principal strains vector) ranged from 48% to 54% (see Table 1, nonlinear case) using the nonlinear model defined in Appendix A. Small force magnitudes were defined as 1/10 of the large force magnitude. Intermediate force magnitudes were defined as the mean value of the large and small force magnitudes (55% of large force). Table 1 also shows maximum strains and averaged maximum strains achieved in the different protrusions of the cells. Fig. 4 shows the ground truth strain measure indicator, i.e. the Frobenius norm of the logarithmic strain tensor on the boundary of the selected cells, for the different cases of considered cellular pulling force levels (see Table 1), assuming material nonlinearity for the hydrogel. It can be observed that the highest levels of strain are concentrated along the protrusion regions of the cells where forces are exerted (see Fig. 2). Note that the range of values shown in Fig. 4 has been chosen to provide a proper visual representation of the strain contours over the cells’ surfaces, and they do not correspond to the maximum values.

**Table 1:** Overall Frobenius norm of the logarithmic strain tensor (ground truth solution) for the nonlinear hydrogel case, for each protrusion of the selected cells for analysis. The values correspond to the case with the highest level of cellular pulling force magnitude (third column).

|  | Protrusion # | Force [pN] | Average | Maximum | Mean average | Mean max. | Maximum max. |
| --- | --- | --- | --- | --- | --- | --- | --- |
| Spread | 1 | 2.25e3 | 0.1870 | 0.5631 | 0.1592 | 0.4884 | 0.6531 |
|  | 2 |  | 0.1215 | 0.2490 |  |  |  |
|  | 3 |  | 0.1690 | 0.6531 |  |  |  |
| Protrusive | 1 | 0.375e3 | 0.1218 | 0.1838 | 0.1064 | 0.2412 | 0.6476 |
|  | 2 |  | 0.0795 | 0.1048 |  |  |  |
|  | 3 |  | 0.1103 | 0.1803 |  |  |  |
|  | 4 |  | 0.1125 | 0.2103 |  |  |  |
|  | 5 |  | 0.0814 | 0.1187 |  |  |  |
|  | 6 |  | 0.1331 | 0.6476 |  |  |  |
| Spherical | 1 | 0.75e3 | 0.2824 | 0.5607 | 0.2158 | 0.3781 | 0.6570 |
|  | 2 |  | 0.1491 | 0.1955 |  |  |  |
| Star-shaped | 1 | 0.67e3 | 0.1303 | 0.2065 | 0.1334 | 0.2602 | 0.4863 |
|  | 2 |  | 0.1457 | 0.2099 |  |  |  |
|  | 3 |  | 0.1390 | 0.4863 |  |  |  |
|  | 4 |  | 0.1086 | 0.1638 |  |  |  |
|  | 5 |  | 0.1660 | 0.3173 |  |  |  |
|  | 6 |  | 0.1109 | 0.1772 |  |  |  |

**Figure 4:**
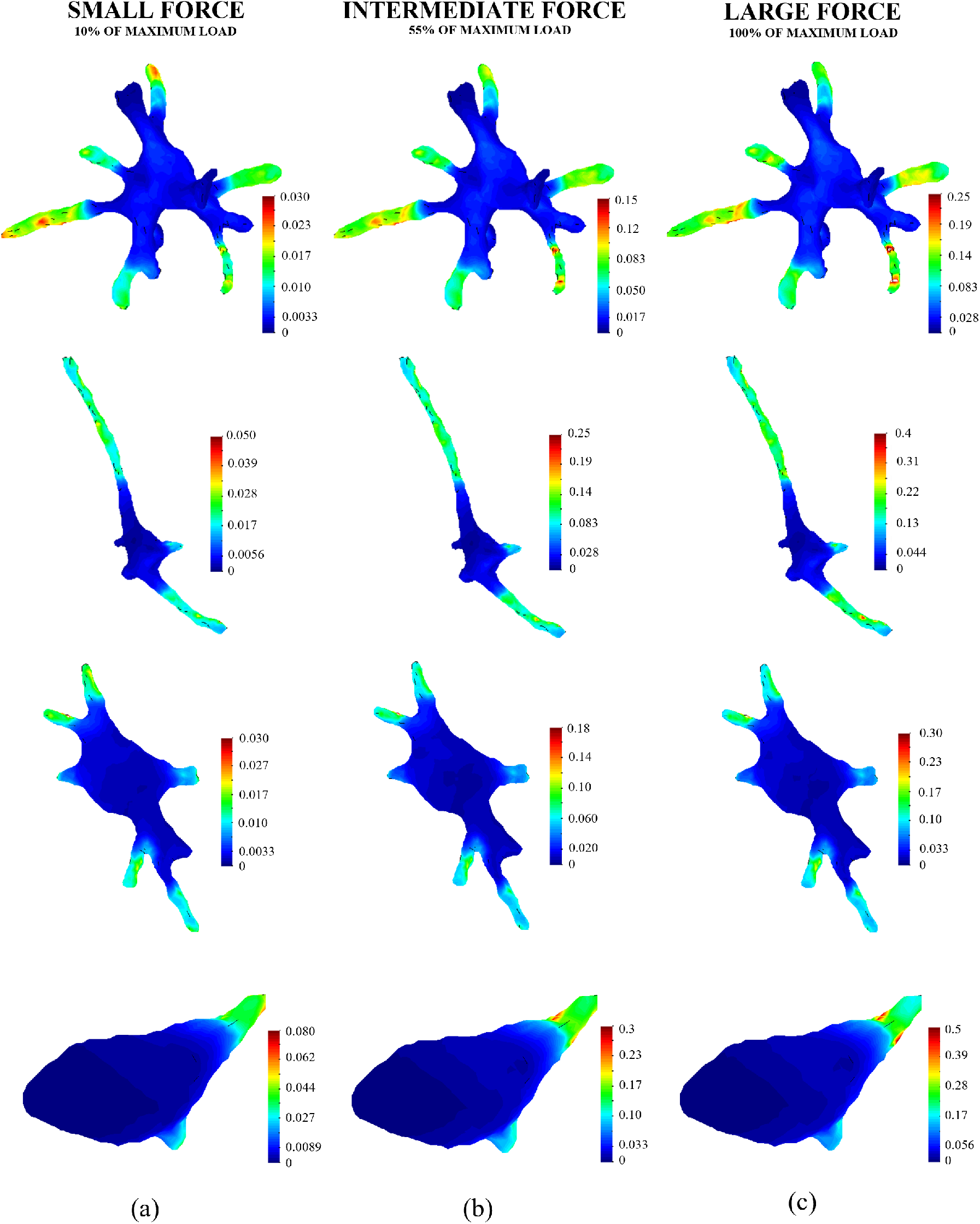
Frobenius norm of the ground truth logarithmic strain tensor [–] on the boundary of the selected cells assuming a nonlinear matrix. (a) small cellular pulling force case; (b) intermediate cellular pulling force case; (c) large cellular pulling force case.

#### Hydrogel’s constitutive behavior

We selected a fibered nonlinear model to describe the nonlinear mechanical behavior of hydrogels. This model has been proven as an excellent fitting for collagen matrices in shear rheology tests (see Steinwachs et al. (2016) and Appendix B). For detailed information about the nonlinear model, the reader is referred to appendices A and B.

Collagen matrices of 1.2 mg/ml concentration were prepared and tested in a shear rheology device following the protocol described in Cóndor et al. (2017). Fig. 5 shows the experimental results of shear rheology tests, as well as the best fit for the selected nonlinear model (see Appendix B). It can be observed, that the model accurately represents the real behavior of the hydrogel up to 50% of shear strain. Table 2 contains the values of the fitted nonlinear model parameters, as well as the parameters fitted for a linear isotropic model. In the latter case, Young’s modulus was computed from the combination of Poisson’s ratio and the shear modulus through the expression 2*G*(1+*ν*). In the literature, one can find Poisson’s ratio values for collagen hydrogels ranging from 0.2 to 0.48 (see Bloom et al. (2008); Bowers et al. (2020); Du et al. (2018); Gjorevski et al. (2015); Koch et al. (2012); Wang et al. (2014)). For this study, 0.34, a value within the range found in the literature, was chosen. The shear modulus was set as the value of the initial slope in the shear rheology experimental curve.

**Table 2:** Fitted nonlinear and linear model parameters. Parameters *E* and *ν* of the nonlinear model stand for the Yong’s modulus and Poisson’s ratio, respectively (see Appendix A for the definition of the nonlinear model parameters).

| Model | Parameter | Value |
| --- | --- | --- |
| Nonlinear | $\kappa_o$ [Pa] | 156.6217 |
| | $d_o$ [-] | 0.375 |
| | $\lambda_s$ [-] | 0.0111 |
| | $d_s$ [-] | 0.0836 |
| Linear | $E$ [Pa] | 29.5076 |
| | $\nu$ [-] | 0.340 |

**Figure 5:**
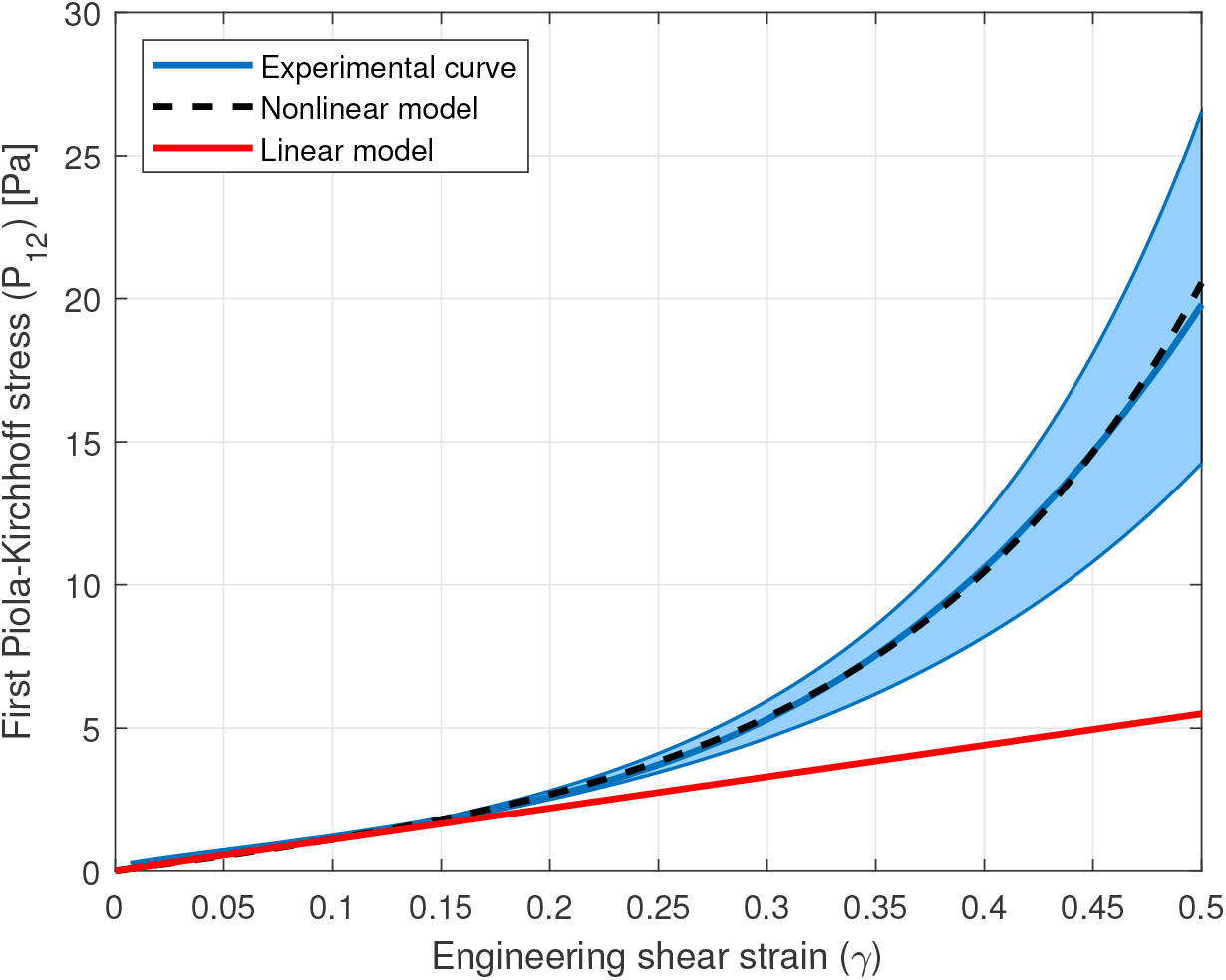
Fitting of experimental curve of 1.2 mg/ml collagen hydrogel with linear elastic and selected nonlinear elastic model for shear rheology tests. The shaded blue area represents the variability of the different tested specimens. The solid blue line curve is the average of all specimens.

The assumed linear and nonlinear models in the different cases of analysis consider the linear and nonlinear fittings shown in Fig. 5. On the one hand, the case of the nonlinear model uses the nonlinear versions of the forward and inverse formulations (finite strain hyperelasticity), as introduced in sections 2.1, 2.2, Appendix A and Sanz-Herrera et al. (2021). On the other hand, the case of the linear model uses the linearized (infinitesimal strains) versions of the forward and inverse formulations (Barrasa-Fano et al., 2021a).

For the different conditions explained above, a total of 24 ground truth displacement fields (4 cells × 3 cellular force levels × 2 material behaviors) were reproduced, see Fig. 1, using the software ABAQUS. These displacement fields, and associated strain and traction variables, are referred to as the ground truth (GT) solutions.

### 3.2. Displacement reconstruction

In a similar way to which it is done in the TFM experimental methodology, GT hydrogel displacements were sampled for each case at discrete random (bead) positions. For all the cases of analysis, we assumed an experimentally feasible bead density of approximately 0.03 beads*/µm*^3^, which relies within the order of magnitude of experimental procedures (Barrasa-Fano et al., 2021a). Table 3 shows the dimensions of the hydrogel domain, the number of beads included in the domain, and the inter-bead distance for each case of analysis. The reconstructed displacement field in the FE mesh is obtained then as a lagrangian interpolation from bead to nodal positions, being the displacements at bead locations previously obtained by interpolation of ground truth displacements present at the nodes in the discretization, whose position does not generally coincide with that of the beads. Fig. 6 shows a schematic representation of this process. We refer to the reconstructed displacement field as the corrupted input displacement field for forward and inverse methods simulations.

**Table 3:** Size of the hydrogel domain, number of beads (*N*_beads_), and inter-bead distance for each cell. The inter-bead distance was calculated as the mean nearest neighbor Euclidean distance.

| | $L_X$ [$\mu m$] | $L_Y$ [$\mu m$] | $L_Z$ [$\mu m$] | $N_{\text{beads}}$ | Interbead distance [$\mu m$] |
| --- | --- | --- | --- | --- | --- |
| Star-shaped | 117.99 | 117.99 | 66 | 28280 | 1.799 |
| Spread | 148.77 | 148.77 | 73 | 49728 | 1.796 |
| Protrusive | 119.13 | 119.13 | 68 | 29703 | 1.802 |
| Spherical | 92.91 | 92.91 | 66 | 17535 | 1.799 |

**Figure 6:**
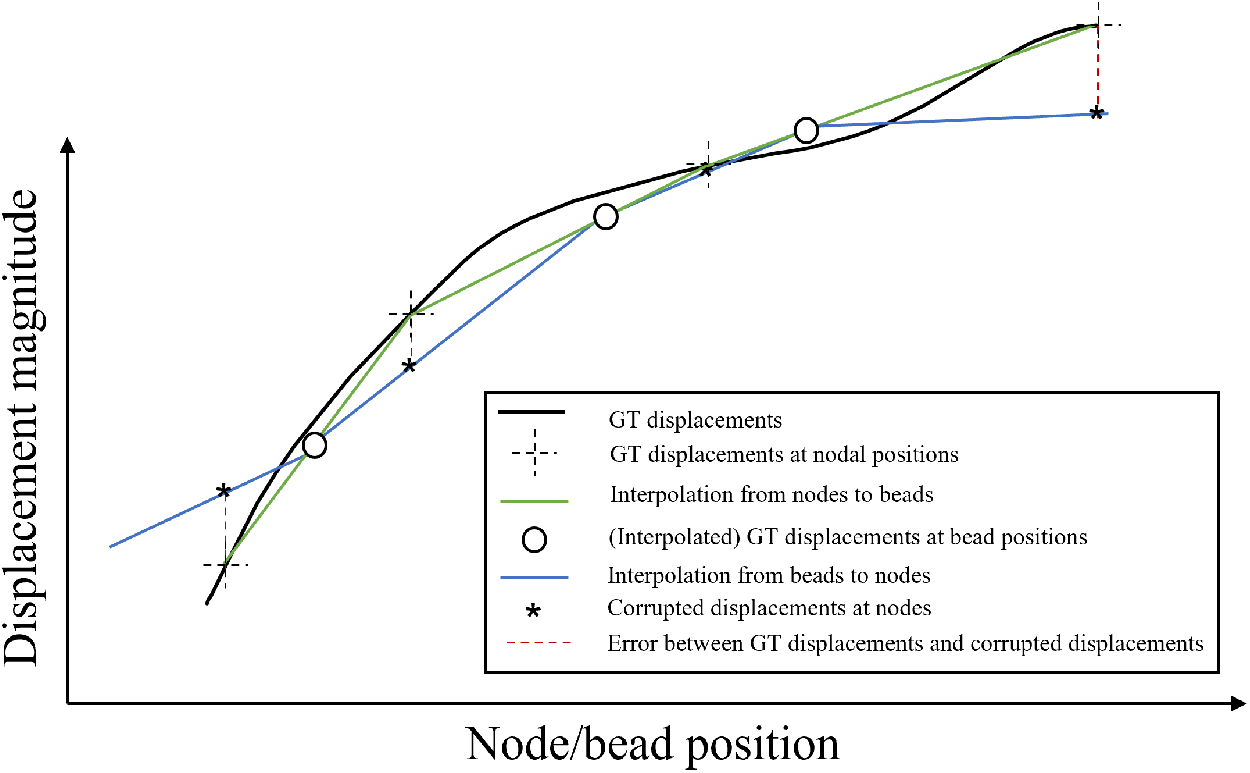
Schematic representation of the process to obtain corrupted displacement fields from ground truth simulations. Errors between ground truth and corrupted displacements were amplified for representation purposes.

With the aim of giving statistical validity to the results, 10 realizations of bead positions for each case were performed corresponding to different instances of the reconstructed displacement fields obtained from the GT (reference) solutions.

### 3.3. Forward and inverse simulations

The different corrupted displacement fields were used as input for the forward and inverse simulations for traction recovery. A total of 120 simulations (4 cells × 3 cellular force levels × 10 realizations) were run for the forward and inverse methods (240 simulations in total) using the linear material behavior, and the recovered displacement field of the linear material case as input. Moreover, another 240 forward/inverse simulations were carried out using the nonlinear material behavior, and the recovered displacement field of the nonlinear material case as input. And finally, a total of 240 forward/inverse simulations using the linear material behavior, and the recovered displacement field of the nonlinear material case as input, were performed. The latter serves to analyze the feasibility of assuming a linear elastic approach (or equivalently neglecting nonlinear effects) for traction reconstruction in nonlinear matrices.

The simulations were performed using ABAQUS in an in-house code orchestrated by Matlab.

### 3.4. Error indicators

In this paper we focus on the errors of traction recovery (either by means of the forward or inverse methods), as they are the most important variable of analysis in TFM. To this end, we define the forward and inverse traction error indicators 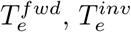 as follows,

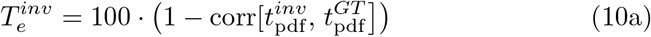

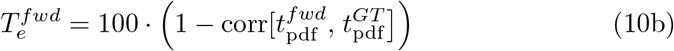

where 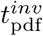 and 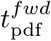 are the probability density functions of the magnitudes of the tractions distribution 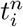 (*i* being the boundary nodes of the cell) for the inverse and forward solutions, respectively. 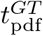 has the same definition but for the ground truth solution. Finally, corr[*f, g*] is the correlation coefficients of functions *f* and *g*.

## 4. Results

The results referred to the different analyses performed in this study (see Fig. 1) are summarized in this section.

The magnitude of tractions on the boundary of the selected cells are represented in Figs. 7–9 for the low cellular pulling force case, including the ground truth solution (either assuming a linear or nonlinear behavior of the hydrogel) and the reconstructed tractions solution for each considered matrix behavior. Specifically, Figs 7 and 8 show the results of reconstructing tractions using the correct (linear and nonlinear, respectively) constitutive behavior (i.e. the behavior used to generate the ground truth, see Fig. 5) by means of the forward and the inverse methods. Finally, Fig. 9 shows the ground truth tractions for a nonlinear matrix, compared to the forward and inverse reconstructed solutions assuming a linear one. Qualitatively, a better performance of the inverse methodology versus the forward one, can be seen in Figs. 7–9 when compared with their corresponding ground truth solutions.

**Figure 7:**
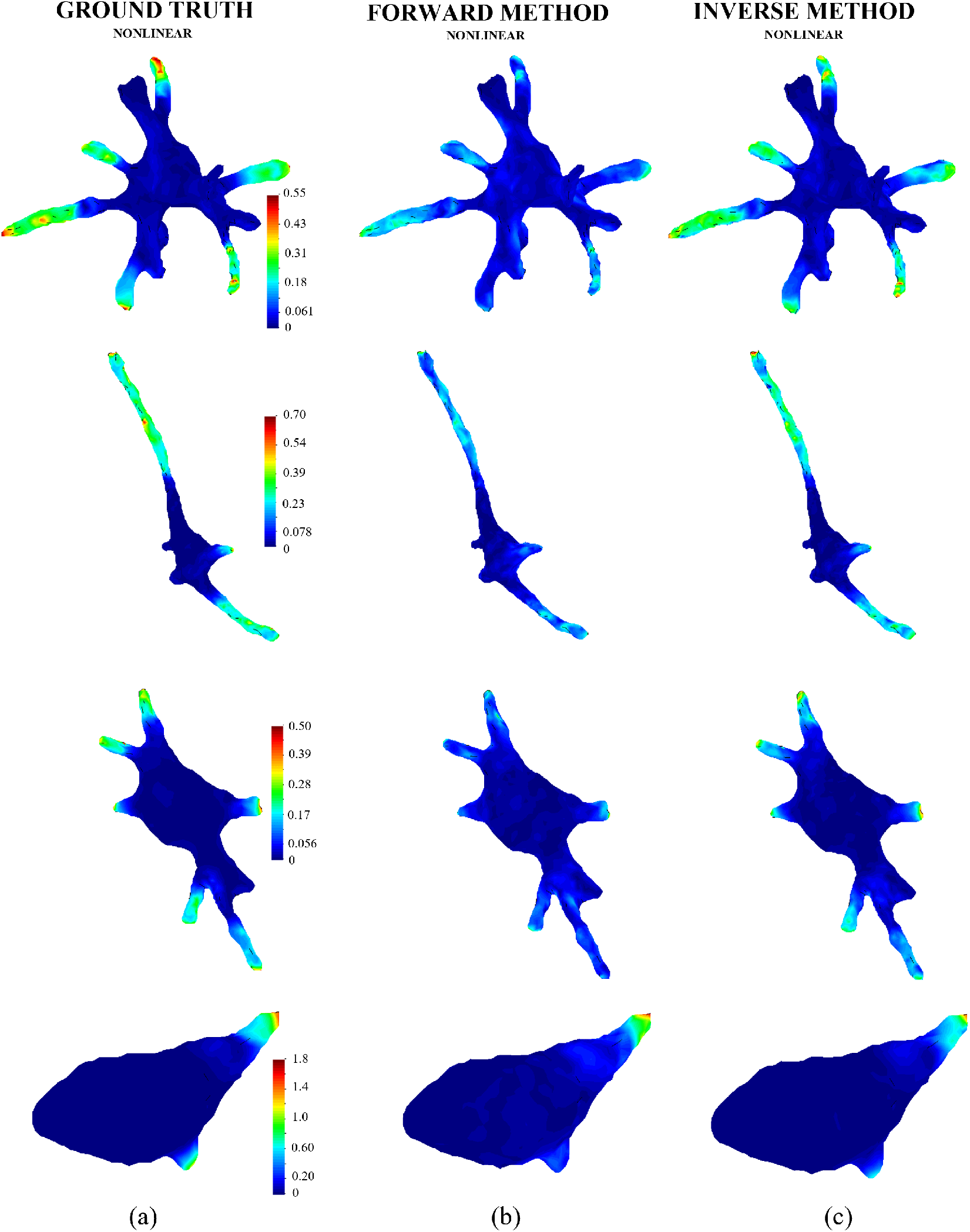
Traction magnitude [Pa] on the boundary of the selected cells. (a) synthetically generated ground truth solution (small cellular pulling force case) in a nonlinear matrix (see Fig. 5); (b) reconstructed solution using the forward method assuming a nonlinear behavior (see Fig. 5) of the matrix; (c) reconstructed solution using the inverse method assuming a nonlinear behavior (see Fig. 5) of the matrix.

**Figure 8:**
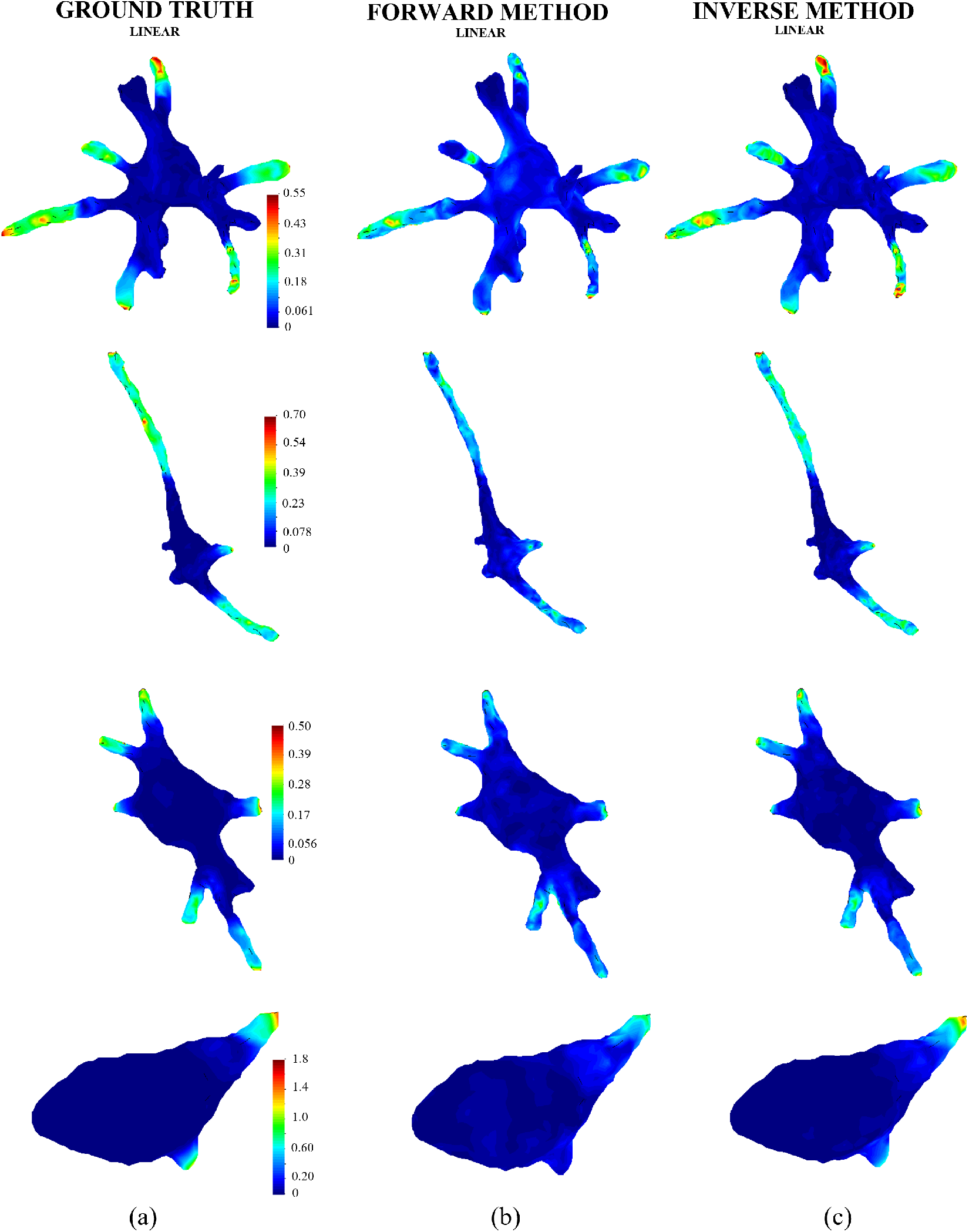
Traction magnitude [Pa] on the boundary of the selected cells. (a) synthetically generated ground truth solution (small cellular pulling force case) in a linear matrix (see Fig. 5); (b) reconstructed solution using the forward method assuming a linear behavior (see Fig. 5) of the matrix; (c) reconstructed solution using the inverse method assuming a linear behavior (see Fig. 5) of the matrix.

**Figure 9:**
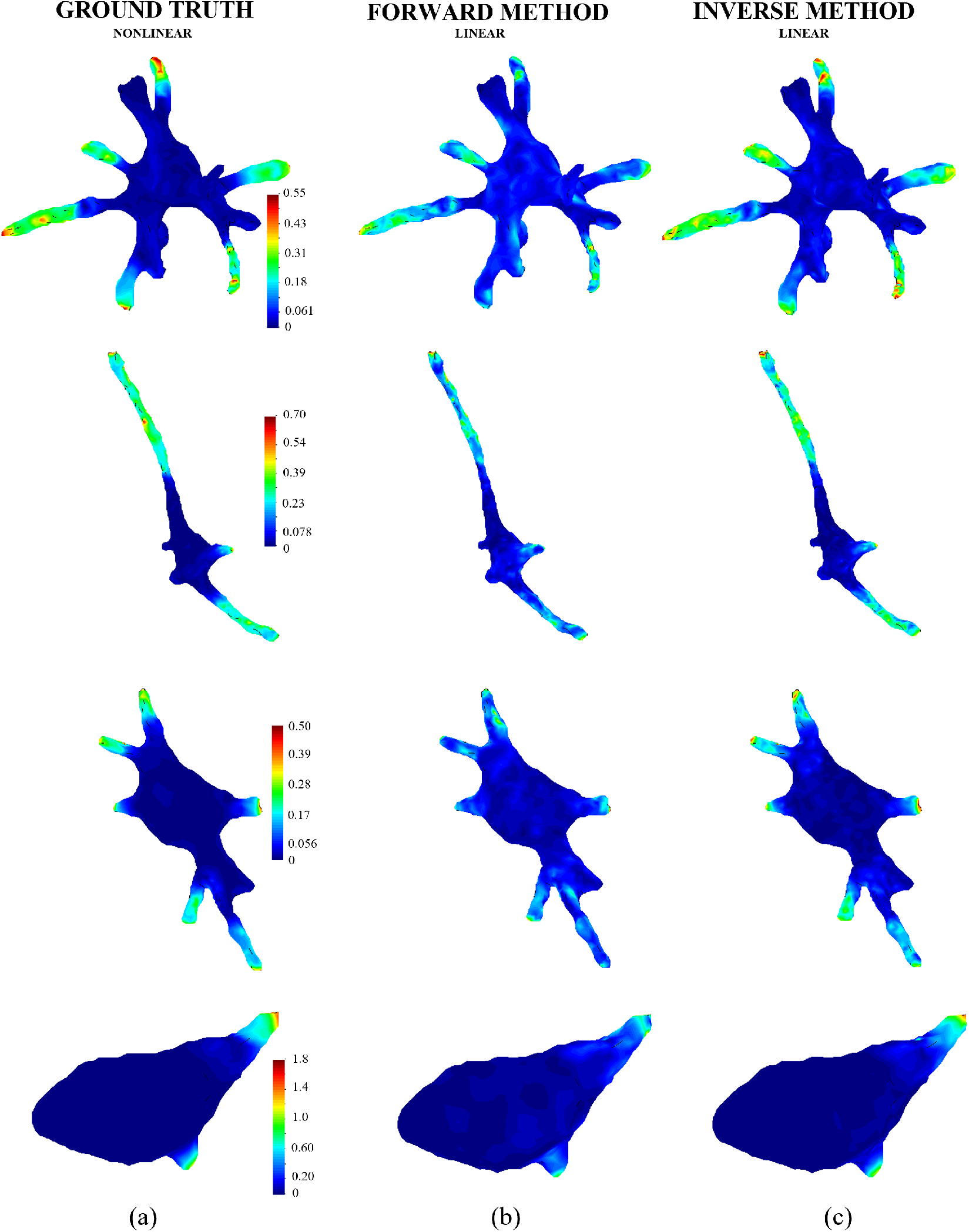
Traction magnitude [Pa] on the boundary of the selected cells. (a) synthetically generated ground truth solution (small cellular pulling force case) in a nonlinear matrix (see Fig. 5); (b) reconstructed solution using the forward method assuming a linear behavior (see Fig. 5) of the matrix; (c) reconstructed solution using the inverse method assuming a linear behavior (see Fig. 5) of the matrix.

Fig. 10 summarizes the errors in the reconstruction of tractions, according to the error indicators defined in Eq. (11), for the different cell morphologies, cellular pulling force levels and methodology (forward and inverse). It quantifies the superior performance of the inverse method for all the analyzed scenarios. The mean values of the traction errors, averaged for the considered pulling force levels, are represented in Fig. 11 versus the solidity index (as a measure of the complexity of the cell geometry) for the forward and inverse methods. Interestingly, it is shown that traction reconstruction becomes more challenging in complex geometries (far from a sphere shape and hence low solidity). A good correlation between the defined traction error indicator and the solidity index was found. Moreover, the outperformance of the inverse versus the forward method is also shown in Fig. 11.

**Figure 10:**
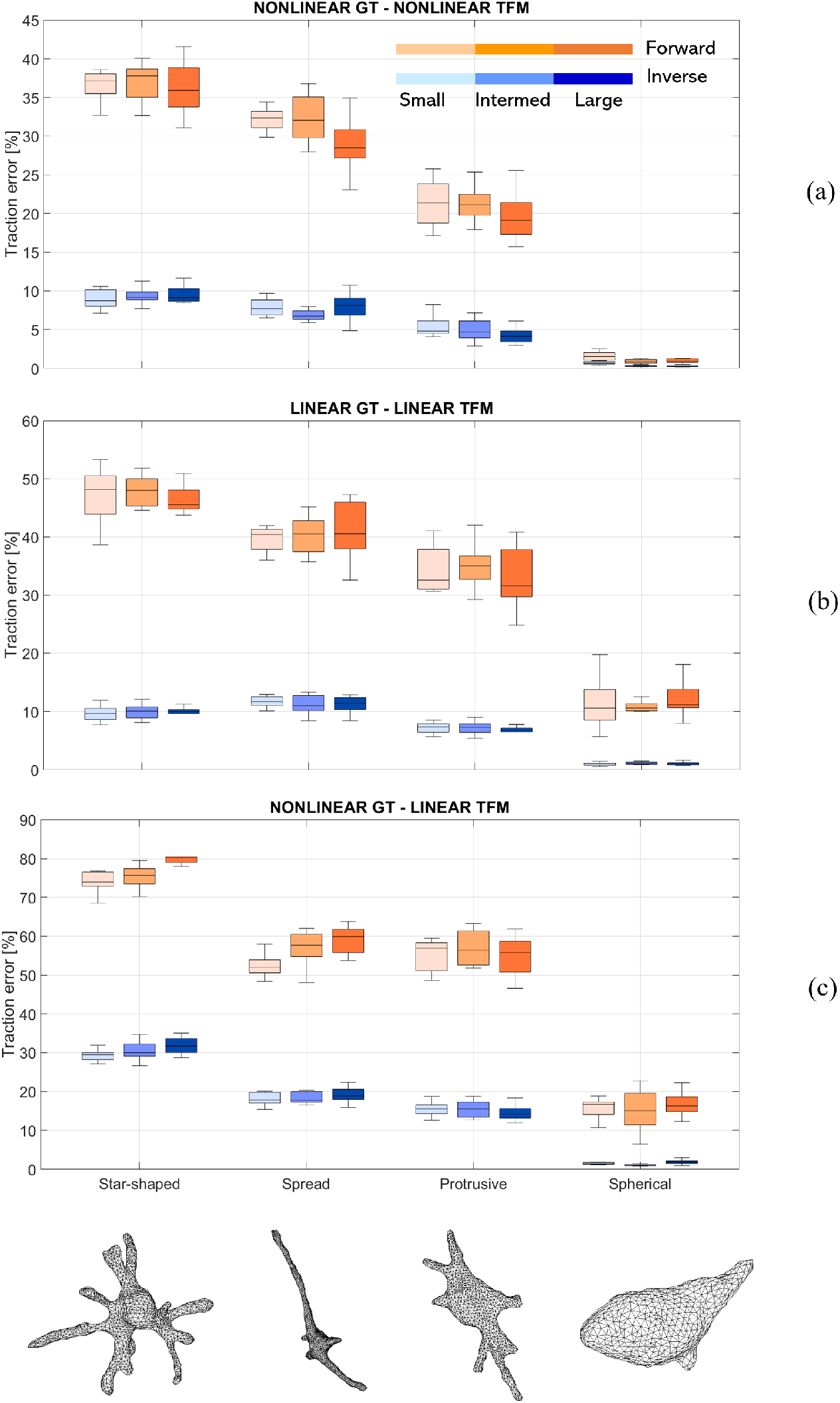
Traction error indicator presented in a boxplot format (10 realizations per considered case) for forward and inverse methods, for small, intermediate and large cellular pulling force cases (see inset legend in plot a). (a) Error of recovered tractions solution assuming a nonlinear matrix versus ground truth solution of the associated nonlinear matrix. (b) Error of recovered tractions solution assuming a linear matrix versus ground truth solution of the associated linear matrix. (c) Error of recovered tractions solution assuming a linear matrix versus ground truth solution of the considered nonlinear matrix.

**Figure 11:**
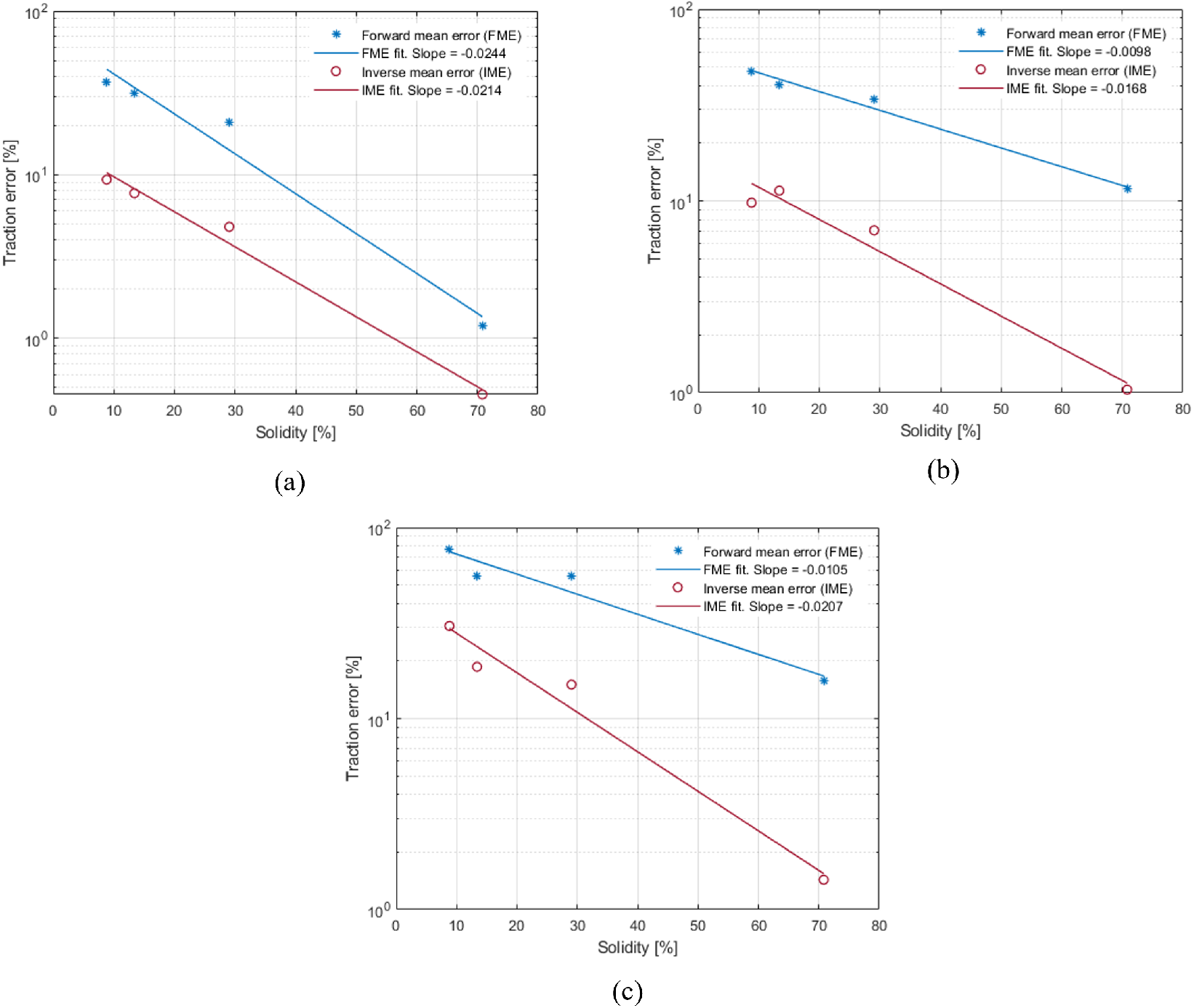
Mean traction error indicator (for analyzed cellular pulling force level cases) of recovered tractions solution: (a) Assuming a nonlinear behavior of the matrix versus ground truth solution of the associated nonlinear matrix. Forward fitting: R^2^= 0.974. Inverse fitting: R^2^= 0.988. (b) Assuming a linear behavior of the matrix versus ground truth solution of the associated linear matrix. Forward fitting: R^2^= 0.986. Inverse fitting: R^2^= 0.969. (c) Assuming a linear behavior of the matrix versus ground truth solution of the considered nonlinear matrix. Forward fitting: R^2^= 0.949. Inverse fitting: R^2^= 0.973.

Mean values of traction errors, CPU time and numerical iterations during the analysis, are represented in Table 4. All the simulations were performed in a standard laptop (AMD Ryzen 7 4800H 2.90 GHz, 16GB RAM). All these values were averaged for the considered pulling force levels and are given for each analyzed cell morphology, hydrogel behavior and methodology (forward and inverse). Moreover, the CPU time is plotted versus traction error (mean values) in Fig. 12 for each considered case. It is observed that the inverse method shows a better accuracy than the forward one, for all the analyzed scenarios, but with a higher CPU time cost. On the other hand, the fact of assuming a linear behavior of the matrix for a nonlinear ground truth behavior, provides a faster but less accurate solution.

**Table 4:** Mean CPU time and mean traction error indicators (averaged over the three analyzed cellular pulling force level cases) of recovered traction solution assuming either linear or nonlinear behaviors of the matrix. Traction errors were calculated with respect to the ground truth solution for the case of the considered nonlinear matrix.

|  | Model | Forward |  | Inverse |  |  |
| --- | --- | --- | --- | --- | --- | --- |
|  |  | Forward time (s) | Forward error (%) | Iters (avg) | Inverse time (s) | Inverse error (%) |
| Spread | Linear | 165 | 55.9 | 1 | 559 | 18.7 |
|  | Nonlinear | 233 | 31.2 | 3.1 | 1849 | 7.7 |
| Protrusive | Linear | 112 | 55.8 | 1 | 234 | 15.1 |
|  | Nonlinear | 156 | 20.7 | 2.7 | 721 | 4.8 |
| Spherical | Linear | 72 | 15.7 | 1 | 116 | 1.4 |
|  | Nonlinear | 96 | 1.2 | 4.3 | 594 | 0.48 |
| Star-shaped | Linear | 124 | 74.7 | 1 | 308 | 30.5 |
|  | Nonlinear | 172 | 36.8 | 3.0 | 993 | 9.3 |

**Figure 12:**
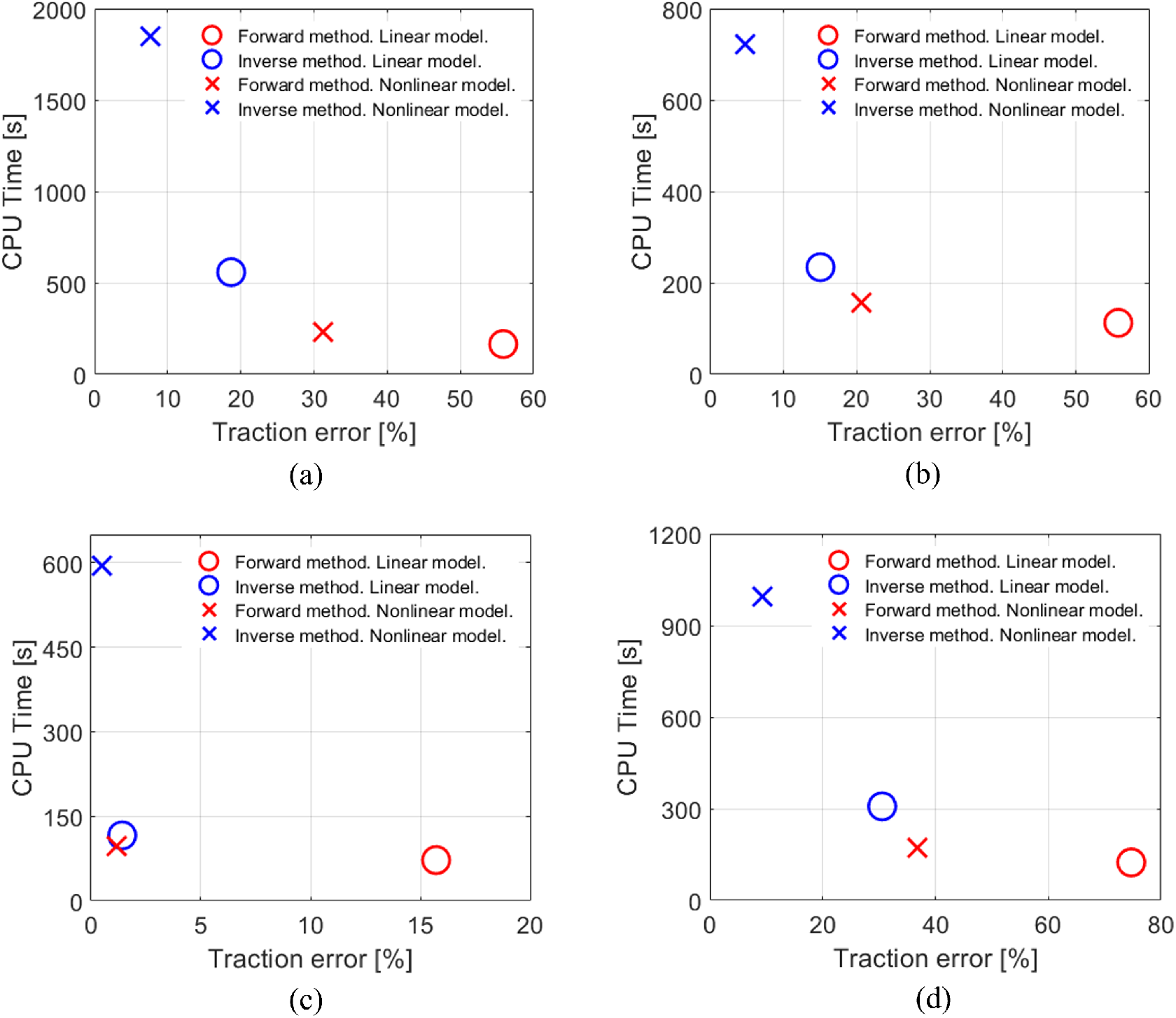
CPU time versus mean traction error indicators (for analyzed cellular pulling force level cases) of recovered tractions solution: (a) Spread cell, (b) protrusive cell, (c) spherical cell, and (d) star-shaped cell.

## 5. Discussion

In this section, we elaborate on the accuracy of traction reconstruction from the different perspectives analyzed in this paper: the magnitude of the cellular pulling force, the effect of the complexity of the cell morphology, accuracy and efficiency of the forward vs the inverse method, and the effect of reconstructing tractions with a correct/incorrect behavior of the hydrogel (linear/nonlinear).

### 5.1. Traction reconstruction accuracy referred to cellular pulling force magnitude

Different cellular pulling force levels were considered, by adjusting nodal forces magnitude in the ground truth simulations, which resulted in different levels of strain in the hydrogel. The magnitudes of the strain measures (Frobenius norm of the logarithmic strain tensor) at the cell–hydrogel boundary ranges from 0.03 to 0.5, for the small and large cellular pulling force cases (see Fig. 4 and Table 1). On the one hand, the case in which a nonlinear hydrogel is considered, the ground truth provides similar strain patterns (but with different magnitude), as represented in Fig. 4, for the different considered pulling force levels. The case in which a linear hydrogel is assumed, the strain patterns are exactly the same but scaled by a certain factor (data not shown). Ground truth tractions on the cell boundary show exactly the same patterns (but scaled) both for the linear and nonlinear hydrogel cases since they are prescribed in the simulations. Therefore, we may assume in a simplified way that the impact of the cellular pulling force in the strain and stress patterns along the boundary of the cells is just a scaling factor in the solution. The traction error for the case in which tractions in a nonlinear hydrogel (nonlinear ground truth solution) are reconstructed assuming a nonlinear behavior of the hydrogel (nonlinear TFM) (Fig. 10a) shows that the impact of the pulling force (and hence strain level) is not substantial on the error of reconstructed tractions, both for the inverse and forward methods. Indeed, the variations of the traction error for the case that tractions in a linear hydrogel (linear ground truth solution) are reconstructed assuming a linear behavior of the hydrogel (linear TFM) (Fig. 10b) are just a consequence of the randomness of the realizations performed. It can be shown along the mathematical formulation of the linear forward/inverse methods (Barrasa-Fano et al., 2021a), that reconstructed tractions are scaled by the same factor that the stress/strain field is scaled in the ground truth simulations. Therefore, the error for a scaled field would remain unaltered for the linear case from a given displacement field.

The case in which tractions in a nonlinear hydrogel (nonlinear ground truth solution) are reconstructed assuming a linear behavior of the hydrogel (linear TFM) requires special attention. In this case, it is considered that tractions are reconstructed using a wrong mechanical behavior of the hydrogel. In addition, nonlinear geometrical effects are also ignored in this case. As a consequence, the largest values of the traction error indicator are produced for this case, as seen in Fig. 10c. Moreover, a larger error with increasing pulling force level would be expected as the assumed geometrical linearity and mechanical behavior of the hydrogel deviates from the true behavior for larger strains (see Fig. 5). Specifically, a slight increasing trend of the error with cellular pulling force level for star-shaped and spread morphologies, but not for protrusive and spherical ones, can be observed. This is a consequence of the definition of the traction error indicator which as a global index that assumes an average of the error for all the tractions (large or small). In order to better explain this observation –as well as others addressed in later sections–, we have defined the ratio between the reconstructed nonlinear and linear strain energy densities, respectively, and checked whether they differ substantially throughout the considered cell surface. Fig. 13 shows the values of the defined ratio for each pulling force magnitude and for each cell geometry. The black regions represent the areas in which the nonlinear strain energy density exceeds the linear one by more than 5% (ratio of 1.05), which is assumed as the large strains scenario where both models present significant differences in their behavior. Although there is an increasing trend in the corresponding magnitude with respect to pulling force magnitude, the black regions constitute no more than a minor fraction of the total surface of the cells. This justifies the observed low sensitivity of the traction error indicator with respect to the pulling force magnitude (Fig. 10c), since it takes into consideration the values of tractions throughout the whole cell surface. It can also be noted that, as seen in Fig. 13, this increase in ratio magnitude is slightly more evident in the case of star-shaped and spread morphologies, whose solidities present the lowest values of the four. Then, the geometrical and material nonlinear effect of cellular pulling force level on the traction error indicator is seen, for this case, when a large pulling force (high strain level) is given through a significant part of the geometry. This situation is found for star-shaped and spread morphologies, according to Fig. 4, but not for protrusive and spherical ones in a substantial way.

**Figure 13:**
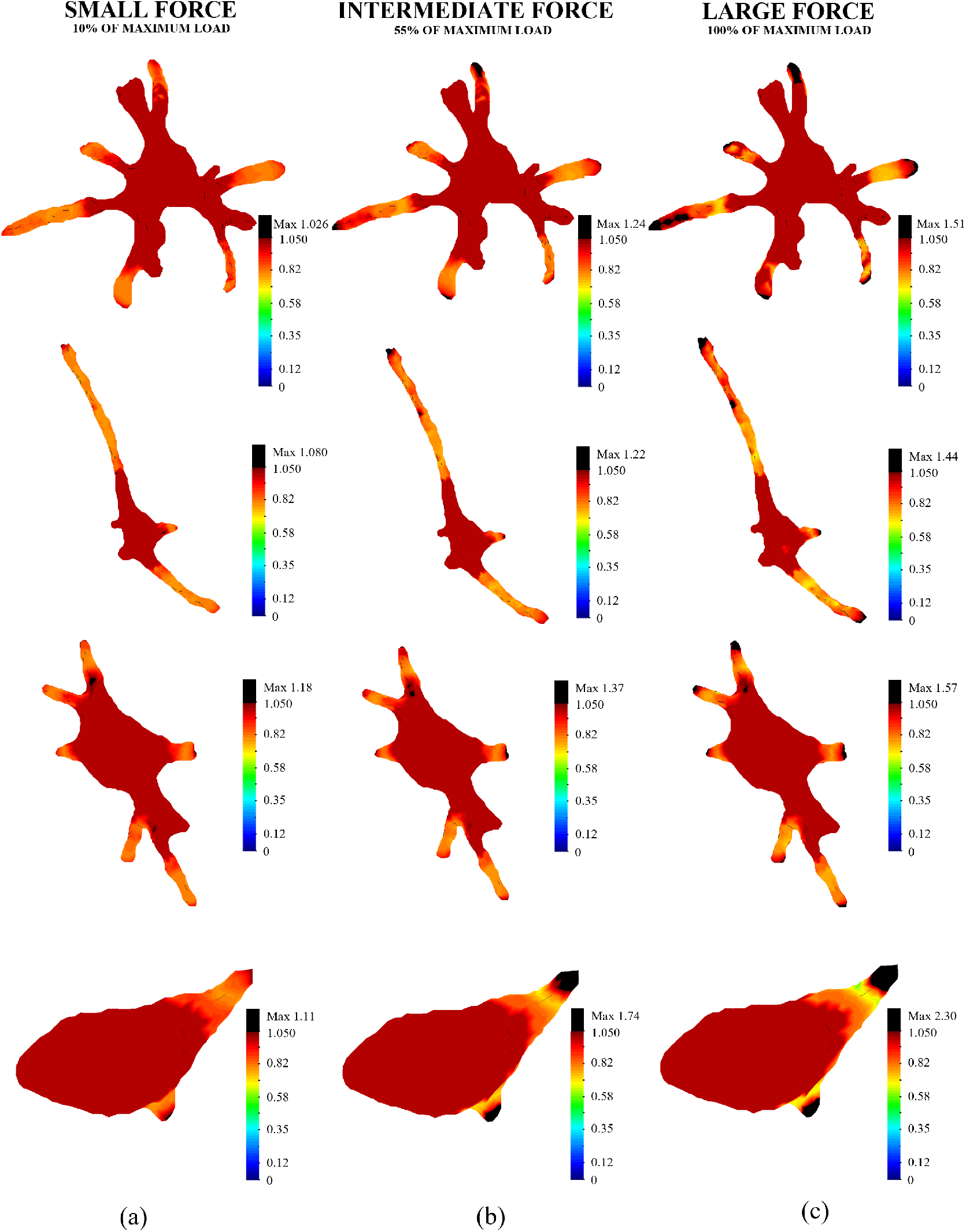
Ratio between reconstructed nonlinear and linear strain energy densities on the boundary of the selected cells assuming a nonlinear hyperelastic behreconstructed solution using the avior for the matrix. (a) small cellular pulling force case; (b) intermediate cellular pulling force case; (c) large cellular pulling force case.

### 5.2. Traction reconstruction accuracy referred to cellular morphology

The impact of cellular morphology on the error of traction reconstruction can be seen in Fig. 11. In this figure, the traction error indicator (averaged for all the pulling force cases) is plotted versus the solidity index of the cell morphology. It is observed that there exists a strong correlation between the error and the solidity index, being more challenging to recover accurate tractions for star-shaped geometries (low solidity indices) than for spherical ones (high solidity indices), for both linear and nonlinear matrices. Indeed, complex geometries (low solidity indices) induce abrupt changes and gradients of displacements in the hydrogel domain near to the cell boundary. Barrasa-Fano et al. (2021a) demonstrated that high gradients of the displacements solution (and hence referred to tractions) are difficult to capture in TFM, as seen here for increasing complexity of the cell morphology.

Furthermore, Fig. 11 shows the error recovery performance as solidity index increases. Besides the superiority of the inverse method, it is shown that the error decreases with a similar trend (in a logarithmic scale) with solidity for the forward and inverse methods, for the case that the nonlinear hydrogel behavior is reconstructed assuming a nonlinear behavior of the hydrogel (Fig. 11a). On the other hand, the error decreases faster (in a logarithmic scale) with solidity for the inverse methods, for the rest of considered cases related to the assumed mechanical behavior of the hydrogel (Figs 11b and 11c).

### 5.3. Traction reconstruction accuracy and computational efficiency referred to forward/inverse methodologies

The outperformance of the inverse method on the accuracy of tractions reconstruction in TFM versus the forward method can be seen in Fig. 10 for all the analyzed cases. Also, these data are plotted in Fig. 11, averaged for all cellular pulling force levels, as it does not represent a significant effect, as discussed before. These figures show errors 3–5 times higher for the forward method compared to the inverse method.

On the contrary, the forward method is computationally less expensive than the inverse method as it includes direct and straightforward algebraic computations, provided the displacement field, either in a linear or nonlinear statement of the problem (Sanz-Herrera et al., 2021). The inverse method requires the setup of a matrix system, and an iterative process in a nonlinear case (Sanz-Herrera et al., 2021). Table 4 and Fig. 12 show the CPU time of the forward and inverse methods, separated for each analyzed case and cell morphology (as it is related to the FE mesh size and hence CPU time). It is observed that higher CPU differences are always found between the forward and inverse for the nonlinear case, as it additionally requires of the mentioned iterative process. The highest CPU time differences are found for the most time consuming case (spread morphology) with a ratio of 233s to 1849s from forward to inverse (for the case that a nonlinear behavior is reconstructed assuming a nonlinear behavior of the matrix), but with a higher traction error cost of 31.2 to 7.7% from forward to inverse, respectively (see Table 4 and Fig. 12).

### 5.4. Traction reconstruction accuracy and computational efficiency referred to hydrogel’s behavior

Linear and nonlinear matrices were considered in the generation of ground truth synthetic solutions, and tractions were reconstructed for both cases considering their corresponding linear or nonlinear behavior. Figs. 10a and 10b show the traction error for nonlinear and linear cases, respectively. It is observed that the errors of the inverse method are similar for all the analyzed cases, whereas the forward provide slightly higher errors for the linear case, specially for the spherical cell morphology. We find an explanation for the last point in Fig. 13. One can observe that there are regions in which the linear strain energy density overcomes the nonlinear one (ratio below 1). This is due to the fact that the considered nonlinear model assumes fiber buckling and hence low stiffness, even negligible, under compression (see Appendix A), whereas the linear model present a constant stiffness under compressive/tensile stresses. Therefore it results in different values that contribute to the error metric, as Fig. 10b shows. These regions are particularly represented in the spherical cell, as illustrated in Fig. 13. Fig. 11 also shows the same trend of the error of the assumed models, but with a faster decrease of the error with solidity index for the linear model (Fig. 11b).

With regards to the CPU time, the linear model runs faster than the nonlinear one for all the analyzed cases, according to Table 4 and Fig. 12. In particular, for the forward method, both the linear and nonlinear methods require only algebraic computations, although with a slightly higher CPU time cost for the nonlinear model since the evaluation of the assumed constitutive law requires a numerical integration over the unit spehere (Steinwachs et al., 2016). Nonetheless, CPU time ratios between the nonlinear and linear model approaches are lower than a 1.4 factor for all the analyzed cases (Table 4 and Fig. 12). On the other hand, the inverse method requires an iterative procedure when the material model is assumed as nonlinear. Despite the few number of iterations until convergence for the inverse nonlinear approach (being 4.3 iterations in average for all the analyzed cellular pulling force levels for the worst case, according to Table 4), CPU times increases to ratios up to 3.3 (worst case), according to Table 4 and Fig. 12, as it increases proportionally to the number of iterations.

#### Feasibility of linear TFM analysis in real nonlinear matrices

The particular case that a nonlinear hydrogel is approached by a linear behavior for traction reconstruction, and hence neglecting nonlinear effects, was analyzed in order to provide an estimation of the feasibility of using linear modeling in TFM even though the real behavior of the matrix fits better with a nonlinear model. A linear model in TFM shows several advantages versus a nonlinear one, such as ease of numerical implementation avoiding convergence troubles, and hence no need for computational mechanics background of the user for troubleshooting; as well as better CPU time performance. Fig. 10c shows the traction error for this case, and Fig. 11c the error averaged for the considered pulling force levels. As expected, it is observed in Fig. 11c that the linear assumption for the hydrogel performs worse in traction reconstruction. Specifically, for the toughest case i.e. the star-shaped morphology, the error increases from 36.8 to 74.7% for the forward, and from 9.3 to 30.5% in the inverse case; when neglecting nonlinear effects in a real nonlinear hydrogel (Figs. 11a and 11c). If we focus on the small cellular pulling force case, such that geometrical nonlinear effects may be neglected, the differences found in Figs 10a and 10c are just a consequence of the differences of the material models used. Despite that linear and nonlinear material models could be assumed as the same for small strains according to Fig. 5, full 3D and multiaxial stress tensors (which is the most frequent stress state in the analyzed cases) show significant differences for the assumed linear and nonlinear models when computed from similar strain fields, even at small strains (specially at compression states as previously commented). On the other hand, CPU time is faster when assuming a linear model instead of a nonlinear one, as discussed before.

## 6. Conclusions

TFM involves many multidisciplinary challenges. From the point of view of mathematical modeling, accurate material models and improved numerical methods are needed to reconstruct reliable tractions, to properly investigate the mechanobiology of in vitro models. In this sense, the impact of a number of issues of interest, such as cell morphology, cellular pulling force and assumed material model, on the traction error recovery; was investigated by means of a conventional forward approach and a version of the inverse method. The inverse method outperforms the forward for all the analyzed cases, reducing the traction error by a factor of 3–5 times. Given the order of magnitude of the traction error, the forward method is not a good approach for the analyzed cases except for spherical morphologies where acceptable errors were found for the forward (∼ 10% for the linear model and < 2% for the nonlinear model).

Regarding the impact of cell morphology on the performance of tractions recovery, it was found a strong correlation between the defined traction error indicator and the sphericity of the cell (solidity index). Moreover, the cellular pulling force level was proven to have a residual impact on the accuracy of traction reconstruction, at least when the error indicator is defined in global terms. The accuracy of traction recovery is similar for linear and nonlinear matrices for the inverse method, but slightly more challenging for linear matrices for the forward method. Despite the advantages that a linear model offers, with better CPU time efficiency amongst other features, it is not a good assumption to represent the behavior of a nonlinear matrix, like collagen hydrogel as used in this study. Even for the inverse method, traction errors range from 15-30% in this scenario.

The knowledge acquired in the analysis performed in this study sheds light on TFM performance in a number of cases of real interest. These situations have not been sufficiently analyzed in previous works. The obtained conclusions may be useful to properly design TFM in vitro experiments, and to further investigate new methods and models in this field.

## Acknowledgements

A.A.-F. and J.A.S.-H. gratefully acknowledge the Financial support from project PGC2018-097257-B-C31 by the Ministerio de Ciencia e Innovación (MCI), Agencia Estatal de Investigación (AEI) and Fondo Europeo de Desarrollo Regional (FEDER); and projects US-1261691 and P20-01195 funded by the Consejería de Economía, Conocimiento, Empresas y Universidad de la Junta de Andalucía. J.B.-F. was supported by KU Leuven internal funding C14/17/111.

## Appendix A

The aim of this appendix is to present a thorough description of the nonlinear model employed in this study (Steinwachs et al., 2016), as well as the derivation process of the corresponding fundamental mechanical relations, to be numerically implemented in the ABAQUS subroutine UMAT, according to the numerical scheme used in this paper. A brief explanation about the Finite Element Method formulation in ABAQUS is given first, in order to provide some context about the variables to be defined in the UMAT subroutine code, to implement a certain constitutive law.

### 4.1. ABAQUS FEM theoretical background

The FE analysis software ABAQUS offers the possibility of implementing any desired constitutive law into its computational process by means of the user subroutine UMAT. This subroutine is coded using the the FORTRAN language.

#### Finite element formulation

The equilibrium of a deformable body can be described through the fundamental scalar expression resulting from the application of the Principle of Virtual Work in spatial configuration,

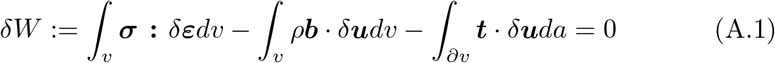

where *δ***u** denotes the virtual displacement, *v* the current volume, *ρ* the current density, **b** the volumetric forces vector, *δϵ* the linear virtual strain tensor, **t** the traction vector resulting from the product between the Cauchy stress tensor *σ* and the vector normal to the surface *∂v*, **n**. A linearized form of Eq. (A.1), in the context of a Newton-Raphson procedure, can be obtained by performing the directional derivative of the virtual work in the direction of the increment Δ**u**. Being *ϕ*_*k*_ a test function,

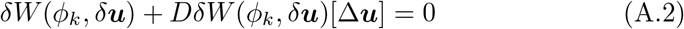

ABAQUS employs the Updated Lagrangian configuration, by means of which the Principle of Virtual Work is expressed as,

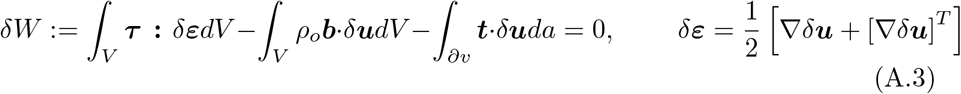

where *τ* denotes the Kirchhoff stress tensor. Note that the volume integrals have been now expressed in the material configuration. At the time of linearizing Eq. (A.3), ABAQUS approximates the term involving the work of internal forces by means of the Jaumann derivative or co-rotational derivative, which represents an objective quantity,

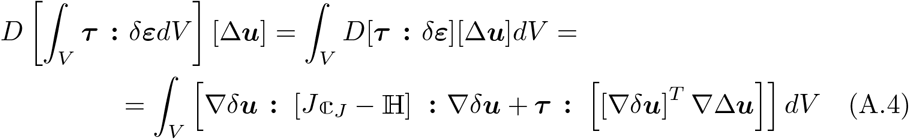

where ℂ^*J*^ is the Jaumann elasticity tensor, which can be computed as,

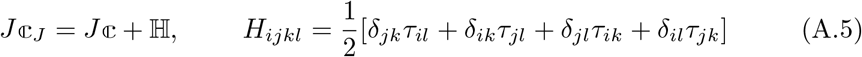

being ℂ the so called Fourth Cauchy Elasticity Tensor,

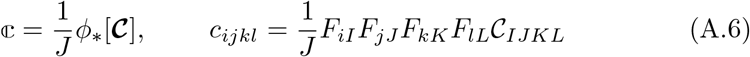

where Einstein’s summation convention has been used.

In order for ABAQUS to accept a certain constitutive relation via the UMAT subroutine, it is necessary to explicitly define both the Cauchy stress tensor and the tangent stiffness matrix DDSDDE as a function of the deformation gradient i.e. *J*ℂ^*J*^. Using the inherent symmetries of both quantities, these need to be expressed in the particular version of the Voigt notation that ABAQUS employs,

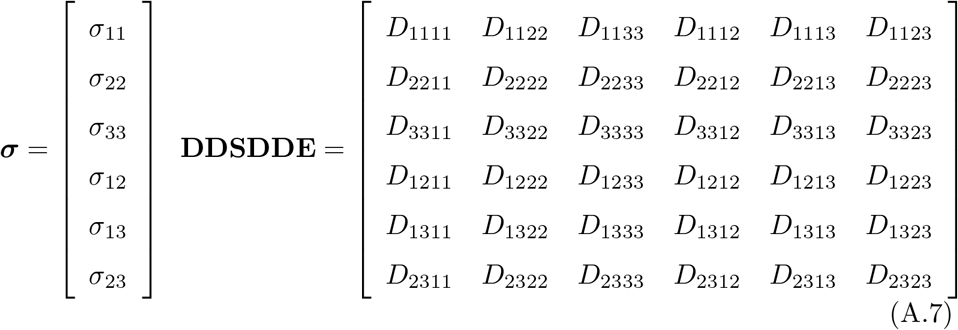

### >A.2. Computational implementation of SAEN model

Steinwachs et al. (2016) propose a model for biopolymer networks based on the combination of a micromechanical basis, i.e., focused on the individual behavior of an individual fiber; and a continuum statement. For a given sufficiently small scale corresponding to a fiber segment, deformations are considered non-affine, this is, independent of deformations of the bulk medium. However, for scales larger tan the typical interconnection distance between fibers, fiber deformations approximate macroscopic deformations *λ*, which depends on fiber orientations and the magnitude of the considered deformations. In this way, deformations are considered to be affine for sufficiently large volumes of the material. The model considers that one fiber can exist in three different states of deformation: compression, in which the fiber becomes unstable, i.e., it buckles, and its stiffness decays exponentially towards zero; straightening, intermediate process in which the fiber recovers its native length after buckling, with constant stiffness; and stretching, a process by which the fiber deforms axially beyond its native length and it experiments an exponential stiffening. The strain energy density function is defined from the stiffness, this is, through its second derivative with respect to the macroscopic deformation measure *λ*,

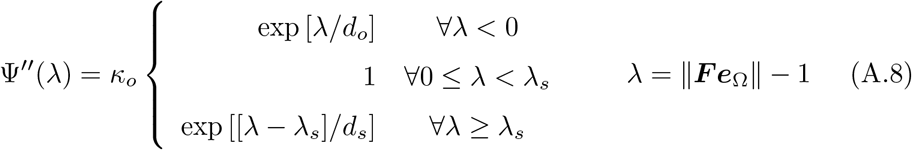

where *κ*_*o*_ is a material parameter with stress dimensions, *d*_*o*_ is the buckling dimensionless parameter, *d*_*s*_ is the strain-stiffening dimensionless parameter and *λ*_*s*_ is the dimensionless parameter that defines the extension of the interval of the linear behavior, i.e., with constant stiffness. The macroscopic deformation *λ* is defined as the variation in length of a fiber with respect its native length, **e**_Ω_ is the unit vector that represents the fiber’s orientation and **F** is the deformation gradient. The mechanical stress of the fiber can be defined by the expression:

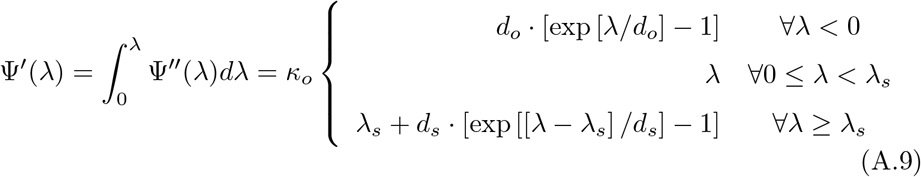

in which integration from zero indicates that the material is not prestressed. The constitutive relation in mixed configuration can then be obtained as,

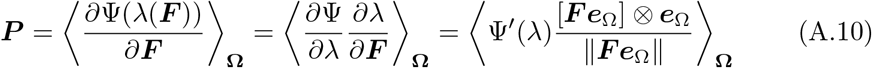

where ***P*** denotes the First Piola-Kirchhoff stress tensor and brackets indicate averaging over all directions Ω in the unit sphere. From Eq. (A.10), the Cauchy stress tensor is obtained by applying a push-forward operation as follows,

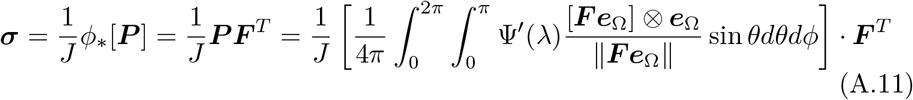

where the integral over the unit sphere is approximated by means of the GaussLegendre quadrature.

The definition of the tangent stiffness matrix DDSDDE depends ultimately on the definition of the Second elasticity tensor ***C***, which can be transformed using Eqs. (A.5) and (A.6) to accommodate to the ABAQUS format. The fourth invariant of the Right Cauchy-Green deformation tensor is described as,

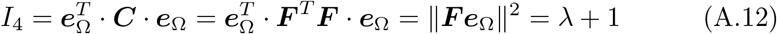

association which permits the rewriting of *λ* in terms of the Right Cauchy-Green deformation tensor,

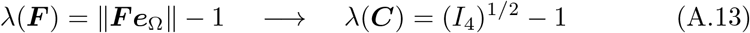

In this way, the constitutive relation in material configuration is obtained as,

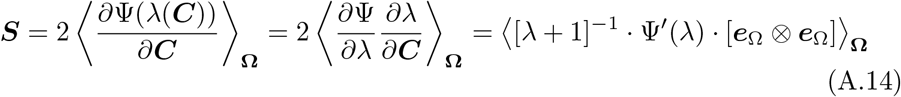

where ⊗ denotes tensor product. The Second elasticity tensor can then be retrieved from,

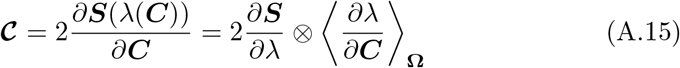

in which,

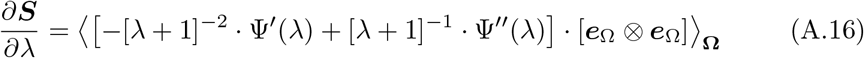

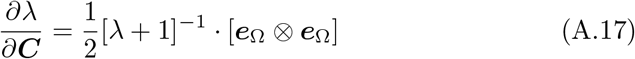

## Appendix B

We fitted the parameters that characterize the SAEN model to experimental rheological data corresponding to a simple shear experiment performed on collagen hydrogels, in order to verify that the model is able to reproduce the real material behavior.

### B.1 Shear rheology

The hydrogel undergoes the action of a rotational rheometer of cone and plate with the aim of measuring the stress-strain relationship within a simple shear state of deformation. The deformation gradient in the case of a simple shear state of deformation is written as,

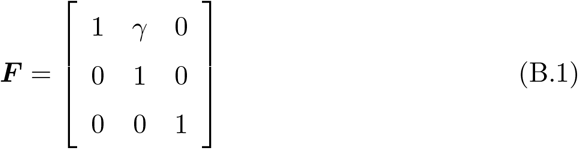

where *γ* represents the engineering shear strain. The shear stress in the test is computed from the measured torque and the setup of the test in the nominal configuration. Therefore, the shear stress resulting from the test corresponds to the shear component of the First Piola-Kirchhoff stress tensor (nominal stress). Recalling the definition given in Eq. (A.10) for this tensor, the nominal stress is then computed from the following expression,

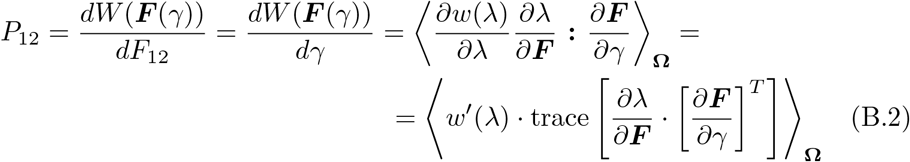

being,

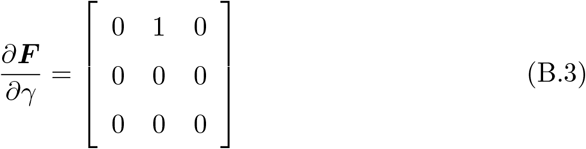

### B.2. Fitting

The fitting of the model parameters have been carried out by means of the MAT-LAB function lsqnonlin, which employs the nonlinear version of least squares minimization. Fig. 5 (main text) shows the fitting performed to the experimental curve corresponding to a hydrogel with a fiber concentrations of 1.2 mg/ml.

## References

Barrasa-Fano, J., Shapeti, A., de Jong, J., Ranga, A., Sanz-Herrera, J.A., Van Oosterwyck, H., 2021a. Advanced in silico validation framework for three-dimensional traction force microscopy and application to an in vitro model of sprouting angiogenesis. Acta Biomaterialia 126, 326–338. doi:10.1016/j.actbio.2021.03.014.

Barrasa-Fano, J., Shapeti, A., Jorge-Peñas, Á., Barzegari, M., Sanz-Herrera, J.A., Van Oosterwyck, H., 2021b. TFMLAB: A MATLAB toolbox for 4D traction force microscopy. SoftwareX 15, 100723. doi:10.1016/j.softx.2021.100723.

Bloom, R.J., George, J.P., Celedon, A., Sun, S.X., Wirtz, D., 2008. Mapping local matrix remodeling induced by a migrating tumor cell using three-dimensional multiple-particle tracking. Biophysical Journal 95, 4077–4088. doi:10.1529/biophysj.108.132738.

Bowers, H.J., Fannin, E.E., Thomas, A., Weis, J.A., 2020. Characterization of multicellular breast tumor spheroids using image data-driven biophysical mathematical modeling. Scientific Reports 10, 11583. URL: https://doi.org/10.1038/s41598-020-68324-4, doi:10.1038/s41598-020-68324-4.

Butler, J.P., Tolić-Nørrelykke, I.M., Fabry, B., Fredberg, J.J., 2002. Traction fields, moments, and strain energy that cells exert on their surroundings. American Journal of Physiology -Cell Physiology 282. URL: http://ajpcell.physiology.org/content/282/3/C595.

Caliari, S.R., Burdick, J.A., 2016. A practical guide to hydrogels for cell culture. Nature Methods 13, 405–414. URL: http://www.nature.com/articles/nmeth.3839, doi:10.1038/nmeth.3839.

Cóndor, M., Steinwachs, J., Mark, C., García-Aznar, J., Fabry, B., 2017. Traction force microscopy in 3-dimensional extracellular matrix networks. Current Protocols in Cell Biology 75, 10.22.1–10.22.20. URL: https://currentprotocols.onlinelibrary.wiley.com/doi/abs/10.1002/cpcb.24, doi:https://doi.org/10.1002/cpcb.24, arXiv:https://currentprotocols.onlinelibrary.wiley.com/doi/pdf/10.1002/cpcb.24.

Dembo, M., Wang, Y.L., 1999. Stresses at the cell-to-substrate interface during locomotion of fibroblasts. Biophysical Journal 76, 2307–2316. URL: http://www.cell.com/biophysj/pdf/S0006-3495(99)77386-8.pdf, doi:10.1016/S0006-3495(99)77386-8.

Discher, D.E., Janmey, P., Wang, Y.L., 2005. Tissue cells feel and respond to the stiffness of their substrate. Science (New York, N.Y.) 310, 1139–43. URL: http://www.ncbi.nlm.nih.gov/pubmed/16293750, doi:10.1126/science.1116995.

Du, Y., Herath, S.C., Wang, Q.G., Asada, H., Chen, P.C., 2018. Determination of Green’s function for three-dimensional traction force reconstruction based on geometry and boundary conditions of cell culture matrices. Acta Biomaterialia 67, 215–228. URL: https://www.sciencedirect.com/science/article/pii/S1742706117307584{#}s0065, doi:10.1016/j.actbio.2017.12.002.

Duval, K., Grover, H., Han, L.H., Mou, Y., Pegoraro, A.F., Fredberg, J., Chen, Z., 2017. Modeling physiological events in 2D vs. 3D cell culture. URL: https://www.ncbi.nlm.nih.gov/pmc/articles/PMC5545611/?report=abstracthttps://www.ncbi.nlm.nih.gov/pmc/articles/PMC5545611/, doi:10.1152/physiol.00036.2016.

Elliott, H., Fischer, R.S., Myers, K.A., Desai, R.A., Gao, L., Chen, C.S., Adelstein, R.S., Waterman, C.M., Danuser, G., 2015. Myosin II controls cellular branching morphogenesis and migration in three dimensions by minimizing cell-surface curvature. Nature Cell Biology 2014 17:2 17, 137–147. URL: https://www.nature.com/articles/ncb3092, doi:10.1038/ncb3092.

Engler, A.J., Sen, S., Sweeney, H.L., Discher, D.E., 2006. Matrix Elasticity Directs Stem Cell Lineage Specification. Cell 126, 677–689. URL: https://www.sciencedirect.com/science/article/pii/S0092867406009615, doi:10.1016/J.CELL.2006.06.044.

Feng, X., Hui, C.Y., 2016. Force sensing using 3D displacement measurements in linear elastic bodies. Computational Mechanics 58, 91–105. URL: https://link.springer.com/article/10.1007/s00466-016-1283-1, doi:10.1007/s00466-016-1283-1.

Franck, C., Maskarinec, S.A., Tirrell, D.A., Ravichandran, G., Genin, G., 2011. Three-Dimensional Traction Force Microscopy: A New Tool for Quantifying Cell-Matrix Interactions. PLoS ONE 6, e17833. URL: http://dx.plos.org/10.1371/journal.pone.0017833, doi:10.1371/journal.pone.0017833.

Gjorevski, N., Piotrowski, A.S., Varner, V.D., Nelson, C.M., 2015. Dynamic tensile forces drive collective cell migration through three-dimensional extracellular matrices. Scientific reports 5, 11458. doi:10.1038/srep11458.

Guilak, F., Cohen, D.M., Estes, B.T., Gimble, J.M., Liedtke, W., Chen, C.S., 2009. Control of Stem Cell Fate by Physical Interactions with the Extracellular Matrix. Cell Stem Cell 5, 17–26. URL: https://www.cell.com/cell-stem-cell/fulltext/S1934-5909{%}2809{%}2900293-8, doi:10.1016/J.STEM.2009.06.016.

Harris, A., Wild, P., Stopak, D., 1980. Silicone rubber substrata: a new wrinkle in the study of cell locomotion. URL: http://www.sciencemag.org/cgi/doi/10.1126/science.6987736, doi:10.1126/science.6987736.

Hervas-Raluy, S., Gomez-Benito, M.J., Borau-Zamora, C., Cóndor, M., Garcia-Aznar, J.M., 2021. A new 3D finite element-based approach for computing cell surface tractions assuming nonlinear conditions. PLoS ONE 16, e0249018. URL: https://dx.plos.org/10.1371/journal.pone.0249018, doi:10.1371/journal.pone.0249018.

Holenstein, C.N., Lendi, C.R., Wili, N., Snedeker, J.G., 2019. Simulation and evaluation of 3D traction force microscopy. Computer Methods in Biomechanics and Biomedical Engineering 22, 853–860. URL: https://www.tandfonline.com/doi/full/10.1080/10255842.2019.1599866, doi:10.1080/10255842.2019.1599866.

Huang, Y., Gompper, G., Sabass, B., 2020. A Bayesian traction force microscopy method with automated denoising in a user-friendly software package. Computer Physics Communications 256, 107313. doi:10.1016/j.cpc.2020.107313, arXiv:2005.01377.

Huebsch, N., Arany, P.R., Mao, A.S., Shvartsman, D., Ali, O.A., Bencherif, S.A., Rivera-Feliciano, J., Mooney, D.J., 2010. Harnessing traction-mediated manipulation of the cell/matrix interface to control stem-cell fate. Nature Materials 9, 518–526. URL: http://www.nature.com/articles/nmat2732, doi:10.1038/nmat2732.

Ingber, D.E., 2002. Mechanical signaling and the cellular response to extracellular matrix in angiogenesis and cardiovascular physiology. URL: http://www.circresaha.org, doi:10.1161/01.RES.0000039537.73816.E5.

Ingber, D.E., 2003. Mechanobiology and diseases of mechanotransduction. URL: http://www.tandfonline.com/doi/full/10.1080/07853890310016333, doi:10.1080/07853890310016333.

Izquierdo-Álvarez, A., Vargas, D.A., Jorge-Peñas, Á., Subramani, R., Vaeyens, M.M., Van Oosterwyck, H., 2019. Spatiotemporal Analyses of Cellular Tractions Describe Subcellular Effect of Substrate Stiffness and Coating. Annals of Biomedical Engineering 47, 624–637. URL: http://link.springer.com/10.1007/s10439-018-02164-2, doi:10.1007/s10439-018-02164-2.

Janmey, P.A., Fletcher, D.A., Reinhart-King, C.A., 2020. Stiffness sensing by cells. Physiological Reviews 100, 695–724. doi:10.1152/physrev.00013.2019.

Jansen, K.A., Donato, D.M., Balcioglu, H.E., Schmidt, T., Danen, E.H., Koenderink, G.H., 2015. A guide to mechanobiology: Where biology and physics meet. doi:10.1016/j.bbamcr.2015.05.007.

Jiang, T., Zhao, J., Yu, S., Mao, Z., Gao, C., Zhu, Y., Mao, C., Zheng, L., 2019. Untangling the response of bone tumor cells and bone forming cells to matrix stiffness and adhesion ligand density by means of hydrogels. Biomaterials 188, 130–143. doi:10.1016/j.biomaterials.2018.10.015.

Kim, S., Uroz, M., Bays, J.L., Chen, C.S., 2021. Harnessing Mechanobiology for Tissue Engineering. Developmental Cell 56, 180–191. URL: https://linkinghub.elsevier.com/retrieve/pii/S1534580720310236, doi:10.1016/j.devcel.2020.12.017.

Koch, T.M., Münster, S., Bonakdar, N., Butler, J.P., Fabry, B., 2012. 3D Traction Forces in Cancer Cell Invasion. PLoS ONE 7, e33476. URL: http://dx.plos.org/10.1371/journal.pone.0033476, doi:10.1371/journal.pone.0033476.

Kopanska, K.S., Alcheikh, Y., Staneva, R., Vignjevic, D., Betz, T., 2016. Tensile Forces Originating from Cancer Spheroids Facilitate Tumor Invasion. PLOS ONE 11, e0156442. URL: https://dx.plos.org/10.1371/journal.pone.0156442, doi:10.1371/journal.pone.0156442.

Kumar, S., Weaver, V.M., 2009. Mechanics, malignancy, and metastasis: The force journey of a tumor cell. Cancer metastasis reviews 28, 113. URL: https://www.ncbi.nlm.nih.gov/pmc/articles/PMC2658728/, doi:10.1007/S10555-008-9173-4.

Kutys, M.L., Chen, C.S., 2016. Forces and mechanotransduction in 3D vascular biology. doi:10.1016/j.ceb.2016.04.011.

Legant, W.R., Miller, J.S., Blakely, B.L., Cohen, D.M., Genin, G.M., Chen, C.S., 2010. Measurement of mechanical tractions exerted by cells in threedimensional matrices. Nature Methods 7, 969–971. URL: http://www.nature.com/articles/nmeth.1531, doi:10.1038/nmeth.1531.

Mammoto, T., Mammoto, A., Ingber, D.E., 2013. Mechanobiology and developmental control. Annual Review of Cell and Developmental Biology 29, 27–61. URL: https://www.annualreviews.org/doi/abs/10.1146/annurev-cellbio-101512-122340, doi:10.1146/annurev-cellbio-101512-122340.

Medina, S.H., Bush, B., Cam, M., Sevcik, E., DelRio, F.W., Nandy, K., Schneider, J.P., 2019. Identification of a mechanogenetic link between substrate stiffness and chemotherapeutic response in breast cancer. Biomaterials 202, 1–11. doi:10.1016/j.biomaterials.2019.02.018.

Nguyen-Ngoc, K.V., Cheung, K.J., Brenot, A., Shamir, E.R., Gray, R.S., Hines, W.C., Yaswen, P., Werb, Z., Ewald, A.J., 2012. ECM microenvironment regulates collective migration and local dissemination in normal and malignant mammary epithelium. Proceedings of the National Academy of Sciences of the United States of America 109, E2595–E2604. URL: https://www.pnas.org/content/109/39/E2595https://www.pnas.org/content/109/39/E2595.abstract, doi:10.1073/pnas.1212834109.

Peng, Y., Chen, Z., Chen, Y., Li, S., Jiang, Y., Yang, H., Wu, C., You, F., Zheng, C., Zhu, J., Tan, Y., Qin, X., Liu, Y., 2019. ROCK isoforms differentially modulate cancer cell motility by mechanosensing the substrate stiffness. Acta Biomaterialia 88, 86–101. doi:10.1016/j.actbio.2019.02.015.

Sabass, B., Gardel, M.L., Waterman, C.M., Schwarz, U.S., 2008. High resolution traction force microscopy based on experimental and computational advances. Biophysical journal 94, 207–20. URL: http://www.ncbi.nlm.nih.gov/pubmed/17827246http://www.pubmedcentral.nih.gov/articlerender.fcgi?artid=PMC2134850, doi:10.1529/biophysj.107.113670.

Sanz-Herrera, J.A., Barrasa-Fano, J., Cóndor, M., Van Oosterwyck, H., 2021. Inverse method based on 3D nonlinear physically constrained minimisation in the framework of traction force microscopy. Soft Matter 17, 10210–10222. URL: https://pubs.rsc.org/en/content/articlehtml/2021/sm/d0sm00789ghttps://pubs.rsc.org/en/content/articlelanding/2021/sm/d0sm00789g, doi:10.1039/d0sm00789g.

Schwarz, U.S., Soiné, J.R.D., 2015. Traction force microscopy on soft elastic substrates: A guide to recent computational advances. Biochimica et Biophysica Acta -Molecular Cell Research 1853, 3095–3104. URL: http://dx.doi.org/10.1016/j.bbamcr.2015.05.028, doi:10.1016/j.bbamcr.2015.05.028, arXiv:1506.02394v1.

Shields, J.D., Emmett, M.S., Dunn, D.B., Joory, K.D., Sage, L.M., Rigby, H., Mortimer, P.S., Orlando, A., Levick, J.R., Bates, D.O., 2007. Chemokine-mediated migration of melanoma cells towards lymphatics -A mechanism contributing to metastasis. Oncogene 26, 2997–3005. URL: https://www.nature.com/articles/1210114, doi:10.1038/sj.onc.1210114.

Song, D., Dong, L., Gupta, M., Li, L., Klaas, O., Loghin, A., Beall, M., Chen, C.S., Oberai, A.A., 2020a. Recovery of tractions exerted by single cells in three-dimensional nonlinear matrices. Journal of Biomechanical Engineering 142. doi:10.1115/1.4046974.

Song, D., Seidl, D.T., Oberai, A.A., 2020b. Three-dimensional traction microscopy accounting for cell-induced matrix degradation. Computer Methods in Applied Mechanics and Engineering 364, 112935. doi:10.1016/j.cma.2020.112935.

Steinwachs, J., Metzner, C., Skodzek, K., Lang, N., Thievessen, I., Mark, C., Münster, S., Aifantis, K.E., Fabry, B., 2016. Three-dimensional force microscopy of cells in biopolymer networks. Nature Methods 13, 171–176. doi:10.1038/nmeth.3685.

Toyjanova, J., Bar-Kochba, E., López-Fagundo, C., Reichner, J., Hoffman-Kim, D., Franck, C., 2014. High resolution, large deformation 3D traction force microscopy. PLoS ONE 9, 1–12. doi:10.1371/journal.pone.0090976.

Vaeyens, M.M., Jorge-Peñas, A., Barrasa-Fano, J., Steuwe, C., Heck, T., Carmeliet, P., Roeffaers, M., Van Oosterwyck, H., 2020. Matrix deformations around angiogenic sprouts correlate to sprout dynamics and suggest pulling activity. Angiogenesis 23, 315–324. URL: http://link.springer.com/10.1007/s10456-020-09708-y, doi:10.1007/s10456-020-09708-y.

Vining, K.H., Mooney, D.J., 2017. Mechanical forces direct stem cell behaviour in development and regeneration. URL: https://www.nature.com/articles/nrm.2017.108, doi:10.1038/nrm.2017.108.

Vitale, G., Preziosi, L., Ambrosi, D., 2012. A numerical method for the inverse problem of cell traction in 3D. Inverse Problems 28. URL: http://iopscience.iop.org/article/10.1088/0266-5611/28/9/095013/pdf, doi:10.1088/0266-5611/28/9/095013.

Vogel, V., Sheetz, M., 2006. Local force and geometry sensing regulate cell functions. URL: http://www.nature.com/reviews/molcellbio, doi:10.1038/nrm1890.

Wang, H., Abhilash, A., Chen, C.S., Wells, R.G., Shenoy, V.B., 2014. Long-Range Force Transmission in Fibrous Matrices Enabled by Tension-Driven Alignment of Fibers. Biophysical Journal 107, 2592–2603. URL: https://www.sciencedirect.com/science/article/pii/S0006349514011096?via{%}3Dihub, doi:10.1016/J.BPJ.2014.09.044.

